# A latent interaction landscape inferred from Micro-C encodes chromatin regulatory identity and conformational diversity beyond contact frequency

**DOI:** 10.64898/2026.03.19.712829

**Authors:** Rahul Mittal, Kavana P. Keshava, Arnab Bhattacherjee

## Abstract

Chromatin contact maps report population-averaged frequencies that obscure the physical principles of genome folding. Here we show that a sparse, signed interaction landscape - a matrix of Lagrange multipliers, ***λ***, inferred by maximum entropy from Micro-C contact probabilities at nucleosome resolution, encodes regulatory and structural information beyond contact statistics. Sign decomposition reveals context-dependent associations with chromatin state. Attractive interactions (***λ***^**−**^) are enriched at active regulatory elements, whereas repulsive interactions (***λ***^**+**^) identify genomic positions whose contacts are depleted relative to the polymer reference, including Polycomb-associated domains. Top-ranked ***λ*** bins show substantially greater enrichment for regulatory chromatin marks than O/E-normalised bins across active, architectural, and repressive classes, demonstrating that ***λ*** preferentially identifies regulatory regions over contact-based rankings. Structural information is hierarchically encoded: the strongest 2% of interactions recover domain boundaries, the top 10% reproduces average contact statistics, yet reconstructing the conformational free-energy landscape, including dominant structural sub-populations and occupancies requires the full interaction spectrum. Nucleosome perturbation experiments show that ***λ*** detects positional information contact maps largely miss, and targeted removal of CTCF/RAD21-associated ***λ*** interactions selectively disrupts loop architecture without altering global polymer statistics. Together, these results establish ***λ*** as a physically interpretable intermediate layer linking chromatin contact maps to three-dimensional structural ensembles.

## Introduction

The three-dimensional organisation of the genome within the nucleus is a fundamental determinant of gene regulation, developmental identity, and cellular function [1–4]. Chromosome conformation capture techniques such as Hi-C and nucleosome-resolution Micro-C have revealed that chromatin folds into hierarchical structures including topologically associating domains (TADs), chromatin loops, and compartmental territories [5–9]. Recent advances in super-resolution imaging and chromatin expansion microscopy have further resolved nanoscale chromatin organisation at the level of individual nucleosomes [10], reinforcing the view that locus-scale architecture is shaped by structural features below conventional Hi-C resolution. Live super-resolution imaging has further revealed that chromatin organises into ∼45-90 nm elongated “blob” domains whose structure and dynamics are spatiotemporally coupled [11]. Yet a fundamental ambiguity persists: experimental contact maps report population-averaged interaction frequencies, not the physical forces or organisational principles that generate them. Many structurally distinct chromatin ensembles can reproduce the same contact map [12–14], and raw contact frequency is dominated by genomic-distance effects that obscure biologically meaningful interaction enrichment. Complementary graph-theoretic and entropy-based approaches have been developed to extract physically interpretable descriptors of multiscale chromatin architecture directly from contact maps [15], yet these remain descriptive rather than inferential. Recovering the physical interaction principles underlying observed contacts therefore requires an inference framework rather than a reconstruction procedure.

A central, yet underexplored, determinant of locus-scale chromatin organisation is nucleosome positioning. Nucleosomes - the fundamental repeating units of chromatin in which ∼147 bp of DNA wrap around a histone octamer - are non-randomly distributed across the genome, with spacing and phasing that vary across promoters, enhancers, and gene bodies [16–18]. This heterogeneous nucleosome-linker architecture modulates local chromatin flexibility, compaction, and the accessibility of regulatory factors including pioneer transcription factors [19, 20]. Crucially, nucleosome positioning varies dynamically across gene bodies and regulatory elements in a transcription-state-dependent manner [21, 22], creating sequence-specific structural heterogeneity that propagates to higher-order chromatin architecture. Recent work has demonstrated that nucleosome positioning alone is sufficient to predict sub-TAD domain boundaries in yeast [23] and that native nucleosomes encode genome organisational principles independently of architectural proteins [20, 24]. Nevertheless, how nucleosome-level heterogeneity quantitatively shapes the effective interaction landscape inferred from high-resolution contact data remains an open question.

Computational approaches to chromatin organisation have largely operated at coarse genomic resolutions of 5-50 kb per bead [25–31], where each polymer unit represents many nucleosomes. While these models have provided important insights into chromosome-scale compartmentalisation and loop organisation, the heterogeneous nucleosome-linker architecture is averaged out by construction. Complementary polymer simulation approaches have shown that heteromorphic chromatin fibre models, in which beads vary in size and compaction according to epigenomic context, can reproduce the three-dimensional folding of complex gene loci [32]. Maximum entropy approaches have proven particularly powerful within this paradigm: MiChroM and related frameworks infer effective interaction energies from Hi-C data at kilobase resolution [26, 33], while the DIMES method [27] uses maximum entropy with pairwise distance constraints from imaging data to generate 3D structural ensembles of interphase chromosomes at kilobase-to-chromosome scale. Complementary Bayesian metainference approaches [34] and experimentally-informed polymer models operating at nucleosome-linker resolution [35] have more recently demonstrated that structural ensembles consistent with Micro-C data can be reconstructed at near-nucleosomal resolution. The challenge of reconstructing three-dimensional genome organisation from contact data spans multiple length scales; early frameworks highlighted the need to bridge the resolution gap between coarse polymer models and the nucleosomal detail encoded in high-resolution contact maps [36]. However, all of these frameworks share a common objective: the primary deliverable is a structural ensemble - a reconstruction of chromatin conformations consistent with the experimental data. The interaction parameters inferred along the way are treated as a means to an end rather than as the object of inquiry.

Here, we adopt a fundamentally different objective. Rather than reconstructing three-dimensional chromatin structures, we ask: *what is the minimal set of effective pairwise interactions required to encode the observed contact statistics?* We develop a maximum entropy framework that infers a sparse matrix of Lagrange multipliers *λ*_*ij*_ - an interaction landscape that encodes the organisational constraints governing locus-scale chromatin architecture - directly from experimental Micro-C contact maps. Chromatin is represented at nucleosome-linker resolution using MNase-seq-derived nucleosome positions [16, 17, 37], introducing sequence-dependent structural heterogeneity into the reference polymer prior to inference. This objective is conceptually distinct from all prior maximum entropy chromatin models: the primary deliverable is not a structural ensemble but the interaction landscape *λ* itself - a compressed, physically interpretable representation that simultaneously corrects for genomic-distance bias, identifies enriched and depleted interactions by sign, and encodes higher-order spatial organisation invisible to raw contact frequency. The sign decomposition of *λ* into attractive (*λ*^−^) and repulsive (*λ*^+^) components provides a mechanistic description of chromatin regulatory state that structural ensembles alone cannot resolve and that distinguishes our approach from all prior maximum entropy chromatin models.

We apply this framework to 12 gene loci in human embryonic stem cells (hESC) and K562 cells, spanning a range of transcriptional states as defined by ChromHMM annotation and RNA-seq expression. We demonstrate that: (i) the inferred *λ* landscape encodes higher-order spatial organisation invisible to contact frequency and hierarchically distributes structural information across the interaction spectrum; (ii) the sign decomposition of *λ* discriminates transcriptionally active from inactive chromatin and outperforms contact-based predictors of regulatory chromatin marks; (iii) controlled nucleosome perturbation experiments reveal that *λ* captures organisational information encoded in nucleosome positioning that contact maps cannot resolve; and (iv) targeted perturbation of CTCF/RAD21-associated *λ* interactions reveals a structured architectural backbone that is biologically organised rather than a generic polymer property. Together, these results establish the inferred interaction landscape as a physically grounded, biologically interpretable layer of chromatin organisation that lies between contact maps and structural ensembles. By treating *λ* as the primary scientific deliverable rather than a means to structural reconstruction, we expose regulatory, architectural, and conformational information encoded in Micro-C data that contact maps alone cannot resolve.

## 1 Results

### The inferred *λ* landscape encodes chromatin interaction principles beyond contact frequency

Experimental contact maps produced by Hi-C and Micro-C report ensemble-averaged interaction frequencies, which is population-level projections of a vast number of structurally distinct three-dimensional chromatin configurations. This many-to-one mapping from conformation to contact is an inherent degeneracy of all chromosome conformation capture experiments [12–14, 38], and it implies that contact maps are necessary but not sufficient to specify the physical interaction principles that organise chromatin. Recovering those principles requires an inference framework that identifies the minimal set of effective interactions consistent with the observed contact statistics.

We address this by casting chromatin organisation as an inverse statistical mechanics problem [28, 39]. Within the maximum entropy (MaxEnt) formulation, a nucleosome-resolved reference polymer ensemble, defined by chain connectivity, bending rigidity, and excluded-volume interactions, is biased by the experimental Micro-C contact probabilities 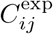. Maximising the relative entropy of the biased distribution with respect to the reference yields the unique least-biased ensemble consistent with the data (Fig. 1A-B):

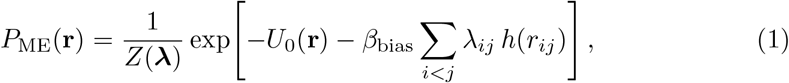

where *U*_0_(**r**) is the reference polymer energy, *h*(*r*_*ij*_) ∈ [0, 1] is a sigmoid contact function encoding spatial proximity between genomic segments *i* and *j*, and *Z*(***λ***) is the partition function. The MaxEnt solution guarantees that 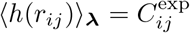 for all constrained pairs, establishing the Lagrange multiplier matrix ***λ*** as the *minimal* set of pairwise couplings required to encode the observed contact structure. Negative *λ*_*ij*_ values represent effective attractions between loci - the interactions occurring more frequently than expected under the reference polymer alone, while positive values represent effective repulsions that suppress proximity below the reference expectation.

**Fig. 1.**
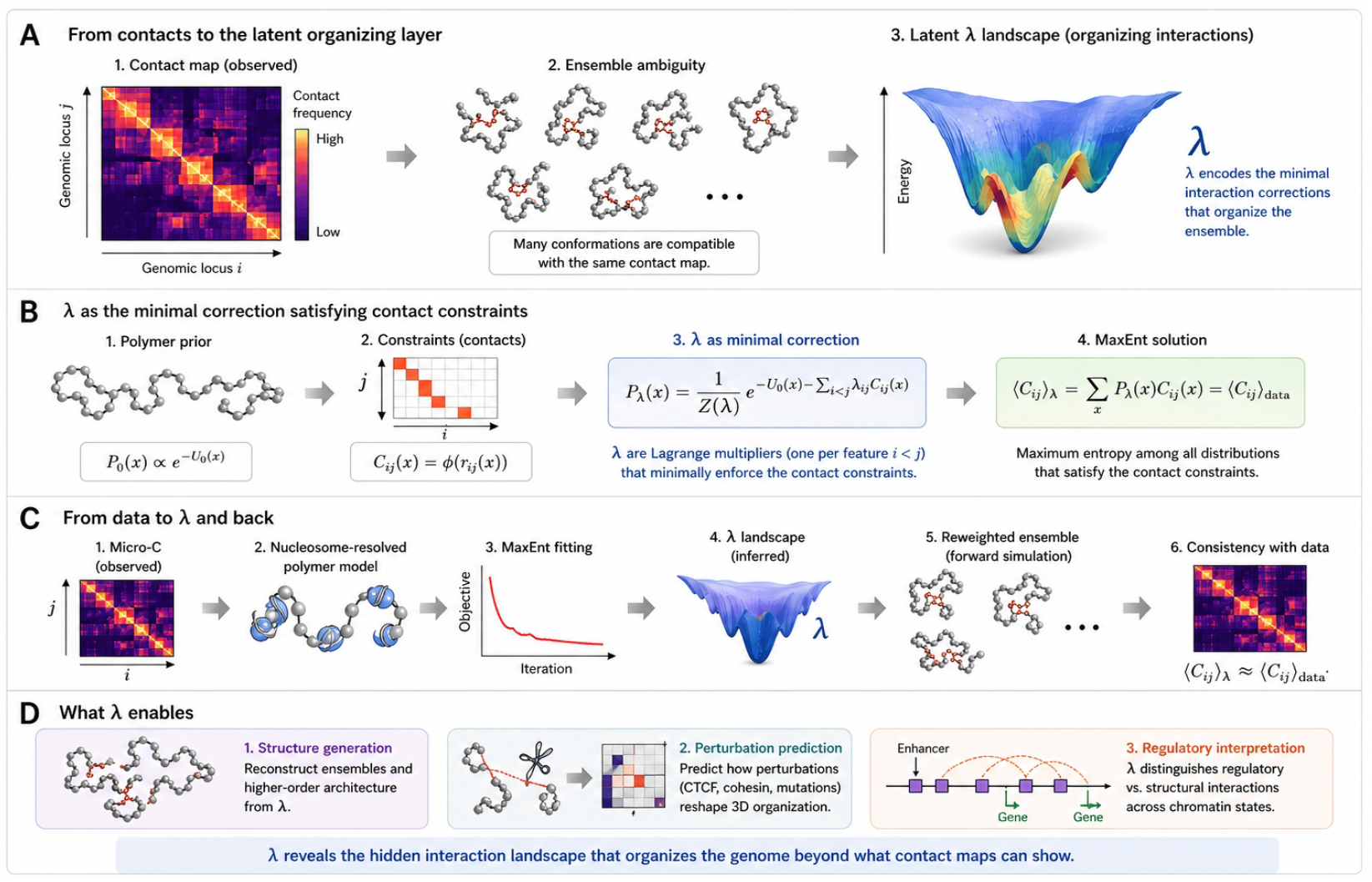
Maximum entropy inference framework and conceptual basis of the *λ* interaction landscape. (**A**) Conceptual motivation. A single experimental contact map is consistent with many structurally distinct chromatin ensembles (left, centre). Maximum entropy (MaxEnt) optimisation identifies the least-biased ensemble consistent with the experimental contact constraints, yielding the interaction landscape *λ*, which specifies the effective interactions required to reproduce the observed contact frequencies (right). (**B**) Mathematical formulation. The reference polymer distribution *P*_0_(**r**), defined by chain connectivity, bending rigidity, and excluded-volume interactions, is biased by the experimental contact constraints *C*^exp^*ij*. MaxEnt optimisation yields the distribution *P* ME(**r**) with Lagrange multipliers *λ*_*ij*_ enforcing ⟨*h*(*r*_*ij*_)⟩_*λ*_ = *C*^exp^*ij*. (**C**) Inference pipeline and validation. Experimental Micro-C data and a nucleosome-resolved polymer model serve as inputs to iterative MaxEnt fitting, producing the converged *λ* landscape. Forward Monte Carlo simulation using the inferred *λ* generates a reweighted structural ensemble whose contact map reproduces the experimental constraints (*C*sim ≈ *C*_exp_). (**D**) Downstream analyses enabled by the *λ* landscape, including structural ensemble generation, predictive perturbation studies, and regulatory interpretation through sign decomposition into attractive (*λ*^−^) and repulsive (*λ*^+^) interaction components.

Three properties make ***λ*** qualitatively distinct from the contact map it is derived from. First, because *λ*_*ij*_ encodes enrichment *relative* to the distance-dependent reference ensemble, it removes the dominant polymer-chain background that inflates contact frequency at short genomic separations, rendering interactions across disparate genomic distances directly comparable. Second, ***λ*** is intrinsically sparse: the vast majority of entries are near zero, and only a small fraction of locus pairs carry significant effective couplings, concentrating the essential interaction signal into a compressed backbone [28, 33]. Third, the sign of *λ*_*ij*_ carries physical meaning inaccessible to contact frequency: two loci may exhibit measurable contacts while being governed by a repulsive *λ*_*ij*_, indicating that their proximity is driven by polymer connectivity rather than by a specific organisational constraint. This sign decomposition provides the basis for the regulatory interpretation developed in the section below on sign decomposition.

The inference pipeline proceeds iteratively from the experimental Micro-C map through MaxEnt optimisation of the nucleosome-resolved polymer model to a converged ***λ*** landscape (Fig. 1C; Methods). Validation by forward Monte Carlo simulation confirms that the inferred ***λ*** reproduces the experimental contact map with high quantitative fidelity (*C*_sim_ ≈ *C*_exp_), establishing that ***λ*** is both necessary and sufficient to encode the observed contact statistics (Fig. 1C). The downstream utility of this landscape is fourfold: it enables full structural ensemble reconstruction, predictive perturbation analysis, regulatory interpretation via the attractive/repulsive sign decomposition, and a unified physical framework linking contact data to functional genomics (Fig. 1D). The following sections characterise each of these capabilities in turn.

### The *λ* landscape defines a sparse, signed interaction landscape beyond distance-normalised contact frequency

A central limitation of raw chromatin contact maps is that contact frequency is dominated by genomic distance. Loci proximal along the linear genome interact more frequently as a direct consequence of polymer chain statistics, irrespective of any locus-specific organisation [5, 12]. This intrinsic bias conflates polymer-driven proximity with genuine interaction enrichment, rendering contact frequency an ambiguous observable. We show that the inferred *λ* matrix resolves this ambiguity and defines a qualitatively distinct interaction landscape, illustrated here for the hESC *nanog* locus.

Distance bias is the first property that *λ* removes. The relationship between |*λ*_*ij*_| and raw contact frequency is distinctly nonlinear (Fig. 2A, left). Although |*λ*_*ij*_| increases monotonically with contact frequency, as reflected in a near-perfect Spearman correlation (*ρ* = 0.996), the substantially lower Pearson correlation (*r* = 0.438) confirms that *λ* is not a linear rescaling of contact data. Data points from widely separated genomic distances fall onto a common monotonic relationship, indicating substantial removal of the dominant distance-dependent bias and rendering interactions across disparate genomic scales directly comparable. Beyond magnitude, *λ* introduces a sign that contact maps cannot carry. Negative *λ*_*ij*_ values correspond to contacts enriched above the distance-expected background, whereas positive values identify pairs depleted below the reference polymer expectation (Fig. 2A, right). This sign decomposition distinguishes interactions enriched above or depleted below the reference polymer expectation - a distinction invisible in any non-negative contact representation. Although O/E normalisation partially corrects for genomic-distance effects, it remains a dense, non-negative transformation of contact frequency that encodes no such signed structure.

**Fig. 2.**
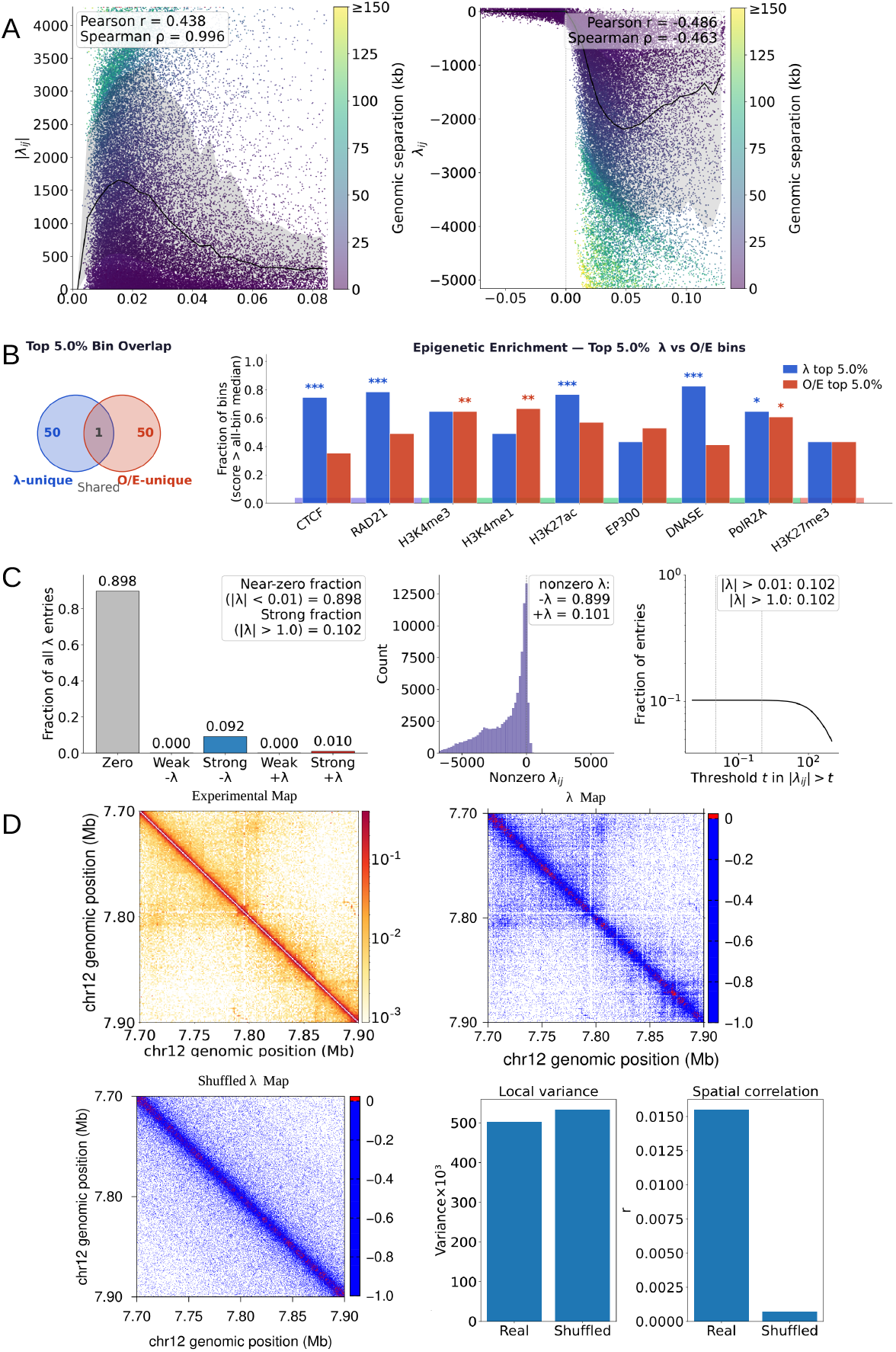
The inferred *λ* interaction landscape is not a transformed contact map, and identifies regulatory genomic elements missed by contact-based representations. (**A**) Left: |*λ*_*ij*_ | versus contact frequency log_10_(1 + *C*_*ij*_), coloured by genomic separation. Near-perfect Spearman (*ρ* = 0.996) and moderate Pearson (*r* = 0.438) correlations confirm a nonlinear, distancedebiased transformation. Right: *λ*_*ij*_ versus residual contact enrichment *C*_*ij*_ −⟨*C*(*s*) ⟩ ; negative values denote contacts enriched above the distance-expected background, positive values denote depleted pairs. (**B**) Left: overlap between the top 5% of genomic bins ranked by |*λ*| and by O/E-normalised contact frequency (hESC *nanog* locus); of 101 bins in each set, only 1 is shared. Right: epigenetic enrichment of *λ*-top-5% (blue) versus O/E-top-5% bins (red) across nine chromatin marks. *λ*-top bins exhibit higher enrichment across all tested chromatin marks than O/E-top bins (Mann-Whitney U-test; asterisks denote p ¡ 0.05-0.001). Results are consistent across three additional loci of contrasting chromatin state (Supplementary Fig. 10). (**C**) Distribution of *λ*_*ij*_ entries: 89.8% near-zero (*λ <* 0.01), with non-zero entries heavy-tailed and predominantly negative (89.9% attractive, 10.1% repulsive). (**D**) Spatial organisation test. Shuffling contacts within genomic-distance bins preserves local variance but collapses spatial autocorrelation to near zero (bottom panels), revealing substantial spatial organisation in the real *λ* map that is absent after distance-preserving shuffling. See Supplementary Fig. 9 for quantitative comparison with O/E. For visualisation of *λmap, λ*_*ij*_*valueswerenormalisedby* max(|*λ*|)*withineachlocusandf orquantitativeanalysesoriginalunnormalisedλvalueswereused*.

A critical question is whether these representational differences have biological consequences. Does *λ* identify genomic elements of regulatory importance that O/E normalisation misses? To address this directly, we compared the top 5% of genomic bins ranked by |*λ*| against the top 5% ranked by O/E-normalised contact frequency (Fig. 2B). The overlap between these two prioritised sets is remarkably small: of 101 bins in each set, only 1 is shared (Fig. 2B, left), establishing that the two representations prioritise almost entirely distinct genomic populations. The *λ*-top bins show consistently higher enrichment across all tested chromatin marks, including active regulatory features (CTCF, RAD21, H3K4me3, H3K4me1, H3K27ac, EP300, DNase hypersensitivity, and POLR2A) and the repressive mark H3K27me3 (Fig. 2B, right; Mann-Whitney *U* -test, *p <* 0.05-0.001). This result indicates that *λ* preferentially prioritises genomic regions enriched for regulatory chromatin features that are not highlighted by contact-based rankings - and that this advantage spans active regulatory, architectural, and repressive chromatin classes, reflecting a broad improvement in biological specificity rather than enrichment for any single epigenomic feature. Consistent enrichment advantages were observed across three additional loci spanning active and inactive chromatin states (hESC *hoxc11*, K562 *nanog*, and K562 *MYC*), with *λ*-top bins outperforming O/E-top bins for the majority of tested marks in every case (Supplementary Fig. 10), confirming that this is a consistent property of the inferred interaction landscape rather than a locus-specific feature.

The interaction landscape that results from this inference is intrinsically sparse. The *λ* matrix carries 89.8% near-zero entries (|*λ*| *<* 0.01), with non-zero entries forming a heavy-tailed distribution strongly skewed toward negative values: 89.9% of non-zero entries are attractive (*λ*^−^) and 10.1% are repulsive (*λ*^+^), with only a small fraction exceeding |*λ*| *>* 1.0 (Fig. 2C). This sparsity directly mirrors the sparsity of the experimental contact map: locus pairs with no detected contact contribute no constraint to the inference and their multipliers remain at zero. Among non-zero entries, the heavy-tailed distribution concentrates organisational signal into a small subset of high-magnitude interactions, suggesting that chromatin architecture may be governed by a limited number of effective constraints rather than the full dense contact matrix - a hypothesis directly tested by the truncation analysis in the following section.

The most stringent test of whether *λ* encodes genuine structural information beyond distance statistics is whether its spatial patterns survive when all distance-dependent marginal statistics are preserved but locus-specific organisation is destroyed. We constructed a null model by permuting contact probabilities within genomic-distance bins, thus preserving the full distance-dependent contact distribution while eliminating locus-specific spatial structure (Fig. 2D). Inference on this shuffled input yields a *λ* landscape visually distinct from the real map. Quantitatively, while local variance is comparable between real and shuffled maps, real *λ* exhibits consistently elevated spatial autocorrelation relative to both the shuffled control and O/E normalisation across all tested lags *δ* (Fig. 2D, bottom right; Supplementary Fig. 9). This dissociation - preserved local variance yet elevated spatial autocorrelation - establishes that the coherent domain-like patterns in *λ* arise from locus-specific long-range dependencies in the contact data, not from marginal distance statistics. Supplementary Fig. 9 further shows that *λ* and O/E define markedly different interaction rankings (*r* = −0.846, *ρ* = −0.816; Supplementary Fig. 9A), consistent with the distinct genomic regions prioritised by the two representations. Critically, the spatial autocorrelation of O/E similarly collapses toward zero across all tested lags, indistinguishable from the shuffled null (Supplementary Fig. 9C), confirming that O/E normalisation does not recover the spatial correlation structure encoded in *λ*.

Together, these results establish that the inferred *λ* matrix is not a normalised representation of contact frequency. Rather, it defines a sparse, signed, and spatially organised interaction landscape that isolates deviations from the polymer background, concentrates structural information into a subset of high-confidence interactions, and exhibits greater specificity for regulatory chromatin features than contact-based rankings.

### A sparse *λ* backbone encodes chromatin domain architecture

Having established that the *λ* matrix captures a structured interaction landscape qualitatively distinct from the contact map, we next examined how architectural information is distributed across the interaction spectrum. Specifically, we asked whether the full *λ* matrix is required to reconstruct chromatin organisation, or whether a sparse subset of high-magnitude interactions suffices to recover the dominant structural features observed experimentally.

Figure 3A contrasts the experimental Micro-C contact map of the hESC *nanog* locus with its inferred *λ* landscape. The inferred *λ* landscape is defined on the same set of experimentally observed locus pairs and preserves the overall sparsity pattern of the contact map, while redistributing interaction strength across those pairs. This redistribution suggests that the physical constraints governing chromatin folding are concentrated within a compact interaction backbone rather than spread uniformly across all locus pairs.

**Fig. 3.**
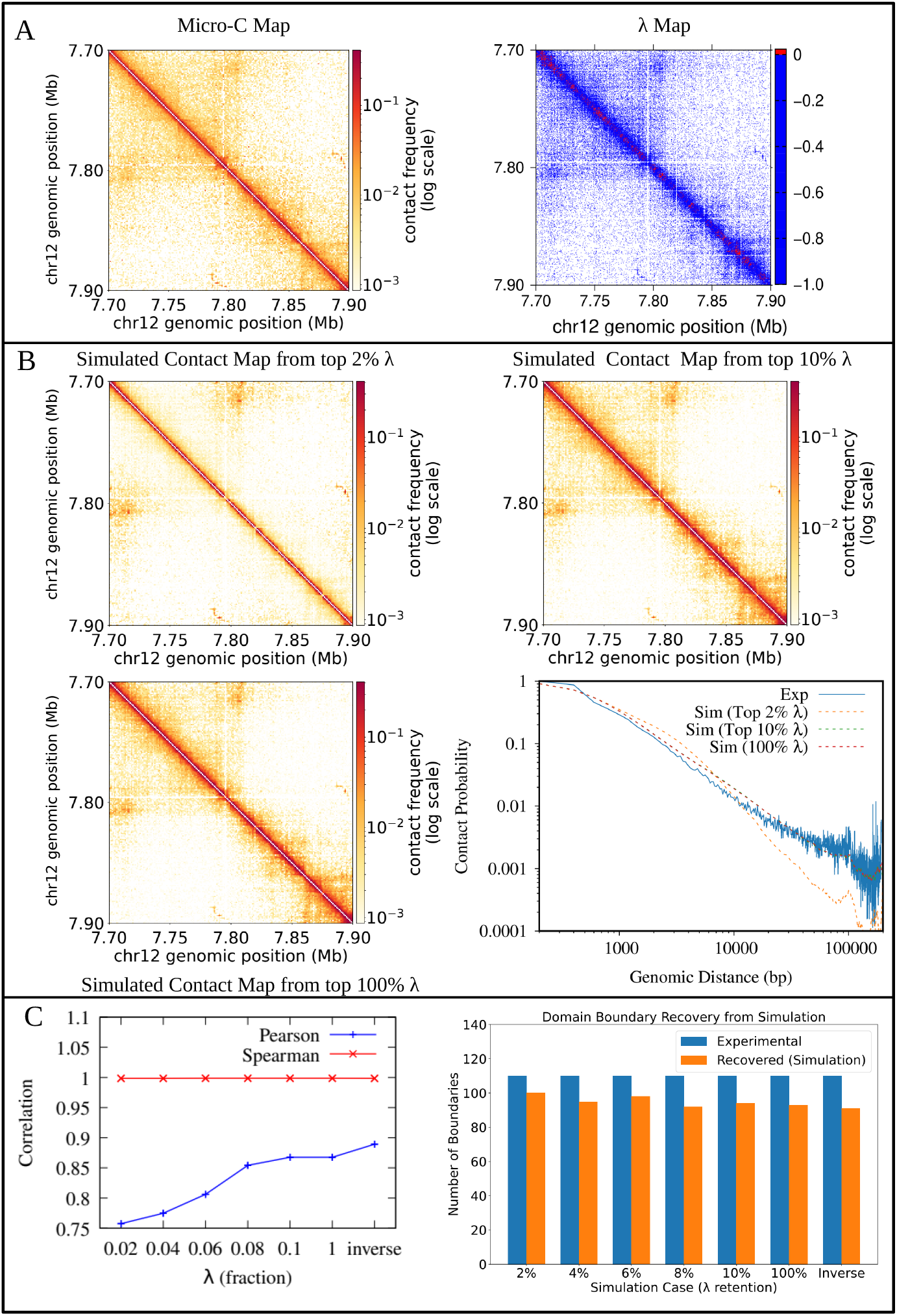
Hierarchical encoding of chromatin structural information within the inferred *λ* landscape. (**A**) Experimental Micro-C contact map (left) and the corresponding inferred *λ* interaction landscape (right) for the hESC *nanog*locus (chr12: 7.70-7.90 Mb). For visualisation of *λmap, λ*_*ij*_*valueswerenormalisedby* max(|*λ*|)*withineachlocusandf orquantitativeanalysesoriginalunnormalisedλvalueswereus(****B*** *simulatedcontactmapsreconstructedf romthestrongest*2%*and*10%*ofinferredλ* interactions (top row) and the full interaction set (bottom left), ranked by absolute magnitude. Contact probability scaling curves *P* (*s*) for all truncation levels versus experiment are shown (bottom right). Intermediate levels (4%, 6%, 8%) are in Supplementary Fig. 11. (**C**) Left: Pearson and Spearman correlations between reconstructed and experimental contact maps as a function of retained *λ* fraction. The Inverse condition denotes a fully iterative MaxEnt simulation in which *λ* is updated until simulated contacts converge to the experimental map. Right: number of insulation-score-defined domain boundaries recovered in simulation relative to experiment across all truncation levels and the Inverse condition.

To test this directly, we ranked all *λ*_*ij*_ entries by absolute magnitude and constructed a series of truncated interaction matrices retaining only the strongest 2%, 4%, 6%, 8%, 10%, and 100% of interactions. Each truncated matrix was used in independent forward Monte Carlo simulations to generate chromatin ensembles and corresponding simulated contact maps (Fig. 3B; intermediate levels in Supplementary Fig. 11). Strikingly, contact maps generated from the top 10% of *λ* interactions are visually nearly indistinguishable from the experimental Micro-C map and faithfully reproduce both domain organisation and long-range contact scaling. By contrast, the top 2% interaction backbone already recovers most domain boundaries and large-scale architectural features, but exhibits noticeable deviations in *P*(*s*) at larger genomic separations, indicating that weaker interactions contribute collectively to quantitative long-range organisation.

Quantitative analysis confirms this picture (Fig. 3C, left). Spearman correlation between simulated and experimental contact maps remains high across truncation levels, indicating preservation of the overall rank ordering of contact probabilities. In contrast, Pearson correlation and contact-probability scaling improve progressively with increasing *λ* fraction, demonstrating that weak interactions contribute primarily to quantitative refinement rather than to the dominant architectural scaffold. The full MaxEnt model achieves the highest Pearson correlation among all conditions, establishing the upper bound on reconstruction fidelity. Remarkably, simulations retaining only the top 2%–10% of *λ* interactions approach this limit in rank-order fidelity, demonstrating that most architectural information is concentrated within a sparse interaction backbone.

We next asked whether this sparse backbone also captures higher-order chromatin organisation. Using insulation score analysis, we identified topological domain boundaries in both experimental and simulated contact maps and quantified boundary recovery at each truncation level (Fig. 3C, right). Even simulations generated from the top 2% of *λ* interactions recover ∼91% of experimentally detected boundaries, and increasing the retained fraction yields only modest further improvement. Domain boundary positions are therefore encoded predominantly within the highest-magnitude tier of the interaction landscape, with weaker interactions contributing to quantitative refinement rather than to the primary architectural scaffold.

Together, these results reveal a hierarchical organisation of information within the inferred *λ* landscape. The strongest few percent of interactions define an architectural backbone that is sufficient to recover most chromatin domain boundaries and the overall topological organisation of the locus. Expanding the retained interaction spectrum progressively improves quantitative fidelity, particularly the long-range contact probability scaling captured by *P*(*s*), indicating that weaker interactions collectively refine chromatin architecture even though they are not required to establish its primary structural scaffold.

### Nucleosome positioning shapes chromatin architecture in ways more strongly resolved by the inferred *λ* landscape than by contact maps

A central question in chromatin biology is whether nucleosome positioning leaves a detectable imprint on higher-order chromatin structure. Population-averaged contact maps may have limited sensitivity to such effects if many distinct nucleosome configurations produce broadly similar ensemble-averaged contact frequencies. We therefore asked whether the inferred *λ* landscape, as a more sensitive reporter of effective interaction constraints, captures structural information that contact maps resolve only weakly or not at all, and whether that information manifests in the properties of the chromatin conformational ensemble.

To address this, we generated three nucleosome-positioning scenarios for the same genomic region of nanog loci (chr12: 7.70-7.90 Mb): (i) experimentally derived positions from MNase-seq, preserving the biologically realistic pattern of irregular spacing, nucleosome clusters, and nucleosome-depleted gaps; (ii) a randomised positioning model in which the same nucleosomes are randomly permuted across the region, preserving nucleosome number but destroying positional organisation; and (iii) a uniform positioning model in which nucleosomes are placed at evenly spaced intervals, eliminating spacing heterogeneity (Fig. 4A). Each positioning model was used to construct an independent reference polymer model, and maximum entropy inference was performed separately to obtain both simulated contact maps and inferred *λ* landscapes.

**Fig. 4.**
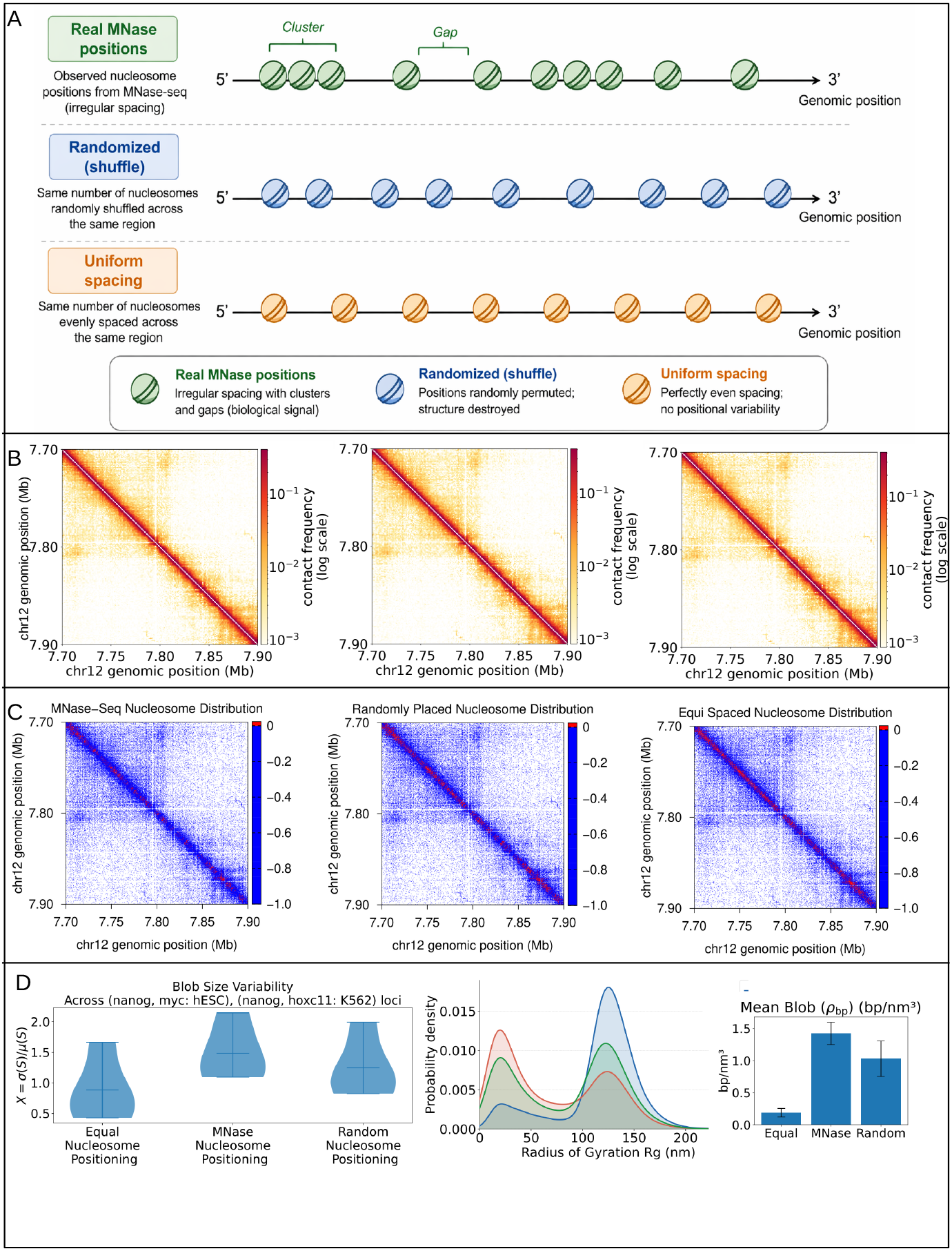
The inferred *λ* landscape is substantially more sensitive to nucleosome positioning than contact frequency maps. (**A**) Three nucleosome configurations used in perturbation experiments: experimentally observed MNase-seq positions (green), randomised positions preserving nucleosome number but destroying positional organisation (blue), and uniformly spaced nucleosomes (orange). All configurations contain the same total number of nucleosomes. (**B**) Simulated contact maps for the hESC *nanog* locus (chr12: 7.70-7.90 Mb) under the three nucleosome configurations. Contact patterns remain broadly similar despite substantial changes in nucleosome organisation. (**C**) Corresponding inferred *λ* interaction landscapes for the same locus. In contrast to the contact maps, nucleosome perturbation produces pronounced reorganisation of the inferred interaction landscape, indicating that *λ* is substantially more sensitive to nucleosome positioning than contact frequency alone. For visualisation of *λmap, λ*_*ij*_*valueswerenormalisedby* max(|*λ*|)*withineachlocusandf orquantitativeanalysesoriginalunnormalisedλvalueswereused*.(**D**)*blob* − *sizeheterogeneityquantifiedbythecoefficientofvariation*X = *σ*(*S*)*/µ*(*S*) across four loci (*nanog* and *myc* in hESC; *nanog* and *hoxc11* in K562). Centre: distributions of blob radius of gyration (*R*_g_) for MNase-derived (green), uniformly spaced (orange), and randomised (red) configurations. Right: mean blob DNA packing density (*ρ*_bp_, bp nm^−3^); error bars denote standard deviation across loci. Together, these measurements indicate that biologically realistic nucleosome positioning promotes heterogeneous chromatin packing characterised by broader blob-size distributions, increased structural heterogeneity, and reduced local packing density, whereas randomisation or uniform spacing yields more homogeneous chromatin organisation.

The simulated contact maps generated under the three nucleosome positioning models are broadly similar (Fig. 4B). The characteristic distance-dependent interaction decay, domain patterns, and near-diagonal enrichment are largely preserved across conditions. Nevertheless, contact maps are not completely insensitive to nucleosome arrangement. A modest but reproducible reduction in Spearman correlation with the experimental map is observed when nucleosome positions are randomised or uniformly spaced, indicating that contact statistics do retain a weak imprint of nucleosome positioning (Supplementary Table 3). However, this sensitivity is limited, and the contact maps do not resolve the structural differences between configurations at a level that would permit discrimination of nucleosome positioning states from contact data alone. This limited sensitivity arises because contact maps represent population averages over large conformational ensembles, in which the spatial heterogeneity introduced by nucleosome positioning is substantially, though not entirely, integrated out.

The inferred *λ* landscapes respond to nucleosome positioning more markedly than the contact maps (Fig. 4C, Supplementary Table 4). Although the global magnitude of *λ* interactions does not change dramatically across configurations, which is consistent with the contact maps being similarly reproduced in each case, the spatial organisation of the *λ* matrix differs systematically and more conspicuously than the corresponding contact map differences. The MNase-derived configuration produces a more structured interaction backbone, with couplings organised into coherent patterns that reflect the underlying nucleosome cluster architecture. The randomised and uniform positioning models yield less organised landscapes, lacking the spatially coherent structure characteristic of biologically positioned nucleosomes. The contrast between *λ* and contact map sensitivity is therefore one of degree rather than kind: both representations carry some information about nucleosome positioning, but the *λ* matrix amplifies and makes explicit a structural signal that remains weak and difficult to interpret in contact frequency space.

Within the structural ensembles generated by the MaxEnt framework, nucleosomes self-organise into spatially compact clusters, hereafter referred to as *nucleosome blobs*. These are locally dense groups of nucleosomes that form the fundamental mesoscale organisational units of chromatin at sub-TAD resolution [35]. These blob-like structures, observed previously in super-resolution imaging of living human cells [11], emerge here directly from the MaxEnt-inferred interaction landscape without explicit prescription of domain boundaries. The most consistent and biologically interpretable signal emerges from analysis of nucleosome blob architecture across configurations (Fig. 4D). The primary finding is that biological nucleosome positioning generates substantially greater heterogeneity in local chromatin domain sizes, as quantified by the coefficient of variation *X* = *σ*(*S*)/*µ*(*S*) of blob sizes, where *S* denotes the size of *i* − th blob and *σ*(*S*) and *µ*(*S*) are standard deviation and mean of all blobs respectively (Fig. 4D, left). This heterogeneity is a robust feature of the configurations produced from MNase-deq positioning data, observed consistently across the nanog and myc loci in hESCs and the nanog and hoxc11 loci in K562 cells, and substantially exceeds the variability produced by the configurations obtained from either the uniform or randomised nucleosome-positioning models. The uniform nucleosome-positioning model, which by construction imposes a homogeneous polymer backbone, produces the most uniform domain sizes, while the randomised nucleosome-positioning model yields intermediate variability. These results indicate that the irregular, non-random spacing of biological nucleosomes, characterised by coordinated clusters and nucleosome-depleted regions, is the primary driver of structural heterogeneity in local chromatin domains.

Complementary to this, the blob *R*_g_ distributions and mean packing density provide additional structural context (Fig. 4D, center and right, Supplementary Fig 12). Individual nucleosome domains in the MNase configuration tend towards smaller radii of gyration and higher local DNA packing density compared with uniform or randomised arrangements. We emphasise that these differences reflect local domain properties - the size and compaction of individual nucleosome blobs - rather than global compaction of the entire locus, which would require independent measurement of locus-wide structural metrics. Within their local spatial extent, however, biological nucleosome clusters assemble into denser and more size-heterogeneous domains than either control configuration produces.

Together, these results reveal a difference in sensitivity between contact maps and the *λ* landscape to nucleosome-scale organisation. Contact frequencies retain only a weak imprint of nucleosome positioning, insufficient to discriminate between biological and non-biological configurations from contact data alone. The *λ* matrix, by contrast, responds more substantially to the spatial organisation of nucleosome positions and faithfully propagates these differences into the structural properties of the inferred chromatin ensemble. The dominant biological signal is not a uniform shift in chromatin compaction but rather an increase in structural heterogeneity. Biological nucleosome positioning generates a diversity of local domain sizes and packing states that is largely absent when positional information is destroyed. This heterogeneity is a hallmark of biological chromatin organisation and is encoded with greater fidelity in the effective interaction landscape than in contact statistics, underscoring the capacity of the *λ* framework to reveal nucleosome-scale structural information that contact maps resolve only weakly.

### The *λ* landscape encodes chromatin regulatory identity beyond contact frequency

Having established that the *λ* matrix encodes a sparse architectural backbone and responds to nucleosome-scale organisation, we next asked whether it also captures biologically interpretable regulatory information beyond that available from contact frequency alone. A key property of the MaxEnt formalism is that *λ*_*ij*_ carries sign: negative values (*λ*^−^) correspond to locus pairs that interact more frequently than expected under the reference polymer ensemble, constituting effective attractive constraints, whereas positive values (*λ*^+^) identify pairs interacting less frequently than expected, constituting effective repulsive constraints. Unlike contact maps, which are strictly non-negative, the *λ* landscape encodes directionality relative to the polymer reference, providing a physically interpretable framework for investigating chromatin-state-dependent deviations from polymer expectation.

We first examined the sign decomposition at two loci with contrasting regulatory identities: the transcriptionally active *ppm1g* locus and the Polycomb-repressed *hoxb1* locus (Fig. 5A,B). At both loci, attractive interactions overwhelmingly dominate the interaction landscape, whereas repulsive interactions constitute a small but spatially structured component. ⟨*λ*^−^⟩*/* ⟨*λ*^+^⟩ = 135.8× at *ppm1g* and 256.1× at *hoxb1*), with repulsive interactions constituting a minor fraction amplified ×10 for visibility. At *ppm1g*, the *λ*^−^ fill and total |*λ*| profile closely track DNase hypersensitivity peaks across the locus, while H3K27me3 is largely absent, consistent with the open, transcriptionally active chromatin state (Fig. 5A). At *hoxb1*, H3K27me3 is the dominant ChIP signal, elevated broadly and continuously across the locus, while DNase signal is sparse; the *λ*^+^ component, though still minor in absolute terms, appears more spatially distributed than at the active locus (Fig. 5B). CTCF shows discrete peaks at both loci without sign preference, consistent with an architectural rather than regulatory function.

**Fig. 5.**
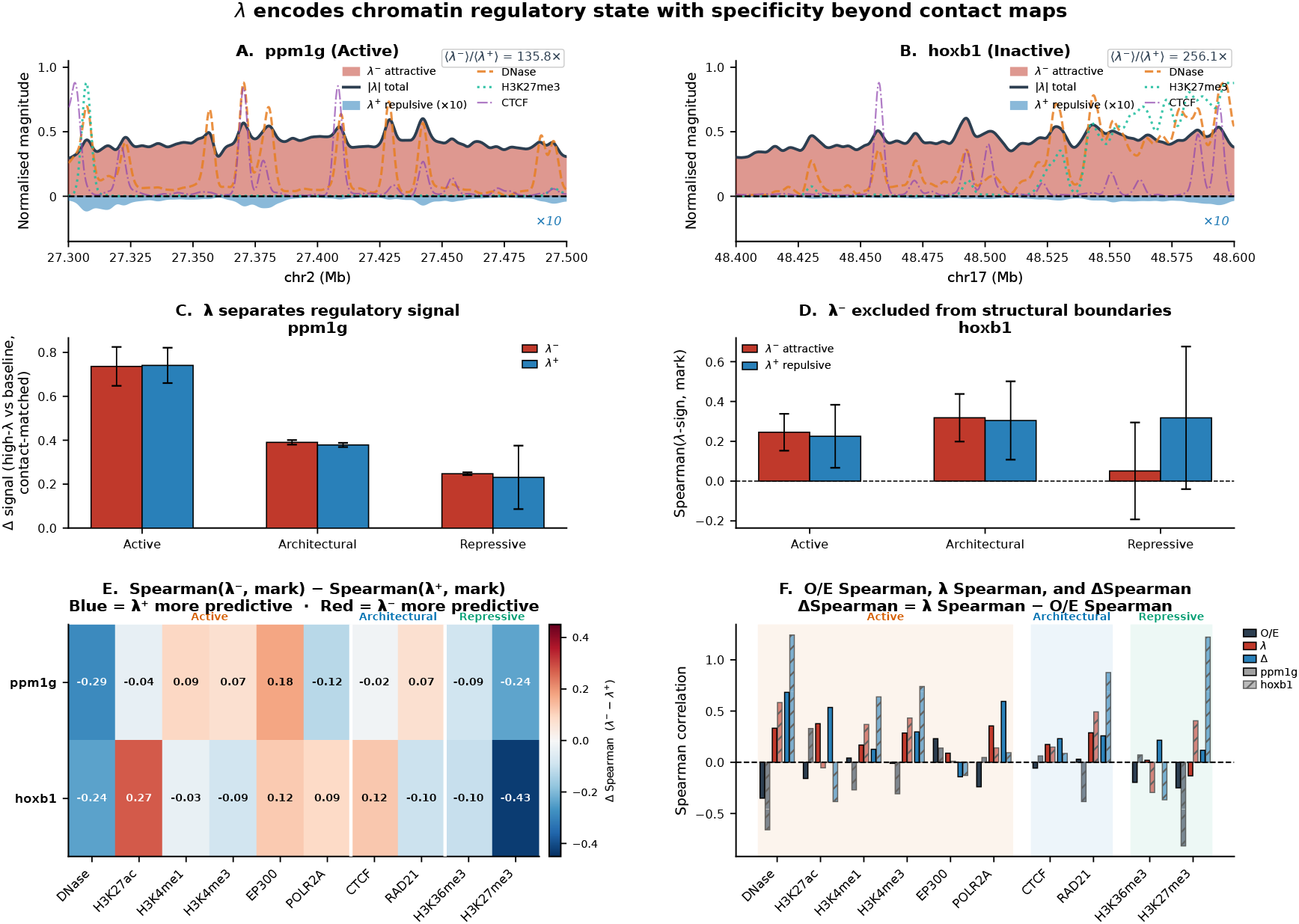
The signed *λ* landscape associates with chromatin regulatory state beyond contact-based representations. (**A, B**) Genomic profiles of attractive interactions (*λ*^−^, red fill), total interaction magnitude (*λ*, black line), and repulsive interactions (*λ*^+^, blue fill, scaled ×10 for visibility), shown alongside DNase hypersensitivity, H3K27me3, and CTCF tracks at the transcriptionally active *ppm1g* locus (chr2: 27.30-27.50 Mb) and the Polycomb-repressed *hoxb1* locus (chr17: 48.40-48.60 Mb). The ratio ⟨*λ*^−^⟩ */* ⟨*λ*^+^⟩ equals 135.8× at *ppm1g* and 256.1× at *hoxb1*.(**C**) Contact-matched enrichment analysis at *ppm1g*. Δsignal quantifies excess regulatory-mark enrichment at high-*λ* bins relative to a contact-frequency-matched baseline, separately for *λ*^−^ (red) and *λ*^+^ (blue), across active, architectural, and repressive mark classes.(**D**) Mean Spearman correlations between *λ*^−^ (red) or *λ*^+^ (blue) and chromatin marks at insulation-defined boundary regions of *hoxb1*, examining sign-specific associations across active, architectural, and repressive mark classes. (**E**) Differential sign-predictivity heatmap showing Δ*ρ* = Spearman(*λ*^−^, mark) − Spearman(*λ*^+^, mark) at *ppm1g* and *hoxb1*. Red indicates stronger association with *λ*^−^; blue indicates stronger association with *λ*^+^. (**F**) Comparison of *λ*-based and O/E-normalised Spearman correlations with chromatin marks at both loci. ΔSpearman = *λ* Spearman − O/E Spearman. For quantitative analyses original unnormalised *λvalueswereused*

A critical question is whether these associations merely reflect information already present in contact frequency, or whether *λ* provides genuinely additional regulatory signal. The contact-matched enrichment analysis at *ppm1g* addresses this directly (Fig. 5C). Genomic bins with high |*λ*| show substantially greater enrichment for regulatory chromatin marks than a background set matched for contact frequency, across active, architectural, and repressive mark classes. Importantly, *λ*^−^ and *λ*^+^ contribute comparably to this excess enrichment - the two bars are virtually indistinguishable across all three mark classes, with fully overlapping error bars. This indicates that it is interaction *magnitude* |*λ*|, rather than sign, that carries contact-independent regulatory information at this locus. The result establishes that the *λ* landscape encodes regulatory signal genuinely beyond what contact frequency alone provides.

We next asked whether the sign decomposition carries positionally specific information at chromatin domain boundaries. Focusing on insulation-defined structural boundary regions at *hoxb1*, we find that active and architectural mark classes show no meaningful difference between *λ*^−^ and *λ*^+^ correlations, with large overlapping error bars in both cases (Fig. 5D). In the repressive mark class, however, *λ*^+^ exhibits higher Spearman correlation than *λ*^−^ at boundary regions, suggesting that repulsive interactions may contribute disproportionately to repressive chromatin organisation at these positions.

The sign-specificity of regulatory associations across individual marks is examined in the differential heatmap (Fig. 5E), which shows Δ*ρ* = Spearman(*λ*^−^, mark) − Spearman(*λ*^+^, mark) at both loci. The pattern is locus-dependent rather than universal. At *hoxb1*, H3K27me3 shows the strongest sign asymmetry with *λ*^+^ substantially more predictive (Δ*ρ* = −0.43), while H3K27ac shows the opposite preference (Δ*ρ* = +0.27), consistent with a partial active/repressive sign dichotomy at this Polycomb-regulated locus. At *ppm1g*, sign preferences are weaker and less consistent: DNase unexpectedly favours *λ*^+^ (Δ*ρ* = −0.29), H3K27me3 also favours *λ*^+^ (Δ*ρ* = −0.24), while EP300 shows modest *λ*^−^ preference (Δ*ρ* = +0.18). A simple active/repressive sign dichotomy therefore does not hold universally, rather the sign-regulatory relationship is most interpretable at loci with strong Polycomb identity.

The overall predictive advantage of *λ* over contact-based representations is quantified in Fig. 5F, which compares Spearman correlations of *λ* and O/E normalised contact frequency with chromatin marks across both loci. *λ* outperforms O/E most clearly and consistently for H3K27me3 at *hoxb1*, where the improvement is the largest in the figure. For active and architectural marks the advantage is generally modest and variable, indicating that the principal gain provided by *λ* arises in chromatin contexts where deviations from the polymer reference are strongest. Both absolute Spearman values and ΔSpearman are reported, confirming that the performance advantage of *λ* is most pronounced where the sign decomposition itself is most informative that is at Polycomb-regulated repressive loci.

The generalisability of these sign-regulatory associations across all 12 analysed loci is examined in Supplementary Fig. 13, which shows Spearman correlations between the *λ*^−^ and *λ*^+^ genomic profiles and chromatin marks across both cell lines and chromatin state classes. Two patterns emerge. First, the most reproducible cross-locus association is a consistently positive correlation between *λ*^+^ and DNase hypersensitivity, which is statistically significant across the majority of loci in both cell lines. Second, positive correlations between *λ*^+^ and H3K27me3 are restricted to Polycomb-regulated loci (*hoxa1, hoxb1, hoxc11*, K562 *tal1*) and are near-zero at active loci. These two patterns together indicate that the biological interpretation of *λ* sign is context-dependent rather than universal. *λ*^+^ does not map simply onto repressive chromatin: instead, the results suggest that *λ*^+^ primarily identifies genomic positions whose contact frequencies fall below the reference polymer expectation, a property that can arise in both accessible and Polycomb-associated chromatin contexts. These observations suggest that *λ*^+^ may arise through multiple biological mechanisms. In accessible chromatin, reduced local compaction could decrease contact frequencies relative to the polymer expectation, whereas in Polycomb-regulated regions compartmental segregation may contribute to similar contact depletion. Distinguishing among these possibilities will require future mechanistic investigation. Conversely, *λ*^−^ shows consistently negative correlation with DNase across loci, indicating that strong attractive interactions preferentially involve genomic positions outside the most accessible chromatin. Together, these cross-locus patterns establish three conclusions. The sign decomposition carries reproducible biological meaning that extends well beyond the two representative loci examined in Fig. 5A-F. The H3K27me3/*λ*^+^ association, while real, is locus-specific rather than a general property of *λ*^+^. And the results suggest that *λ* sign primarily reflects whether contacts are enriched or depleted relative to the polymer reference ensemble, whose chromatin-state correlates depend on which biological context drives that deviation at a given locus.

Together, these analyses indicate that *λ* interaction magnitude captures regulatory information beyond that contained in contact frequency alone. At the Polycomb-regulated hoxb1 locus, *λ*^+^ exhibits its strongest associations with repressive chromatin features, particularly H3K27me3. Whereas contact maps quantify how frequently loci encounter one another, *λ* quantifies whether those encounters occur more or less often than expected from polymer physics alone, a distinction that enriches regulatory resolution beyond what contact frequency encodes.

### The *λ* landscape governs the chromatin free-energy landscape and hidden conformational organisation

The inferred *λ* landscape determines not only the mean contact map of a chromatin locus but also the full free-energy landscape explored by the corresponding conformational ensemble. A central question is whether this landscape is effectively captured by conventional low-dimensional structural observables, or whether it harbours hidden conformational organisation that such descriptors cannot resolve. We addressed this by combining global structural descriptors with contact fingerprint analysis of the hESC *nanog* locus ensemble, and by comparing the effects of distinct perturbations to the interaction landscape.

Projection of the ensemble onto the radius of gyration (*R*_*g*_) and long-range contact fraction (*f*_LR_) reveals a single dominant free-energy basin (Fig. 6A, left), suggesting apparent structural homogeneity at the level of global observables. However, projection of the identical ensemble into Blob-PCA contact fingerprint space produces a qualitatively different picture (Fig. 6A, centre). Multiple distinct conformational substates emerge. For example, some of the dominant ones (*>* 5% occupancy) are S1 (27.5%), S3 (24.4%), and S4 (20.5%). Each of them shows a characteristic spatial organisation despite overlapping *R*_*g*_ and *f*_LR_ values, confirming that global compaction and long-range contact content are insufficient to distinguish between them. Representative conformations extracted from each substate display visually distinct chromatin architectures (Fig. 6A, right), demonstrating that what appears as a single basin in conventional low-dimensional structural observables conceals a mosaic of structurally distinguishable chromatin configurations.

**Fig. 6.**
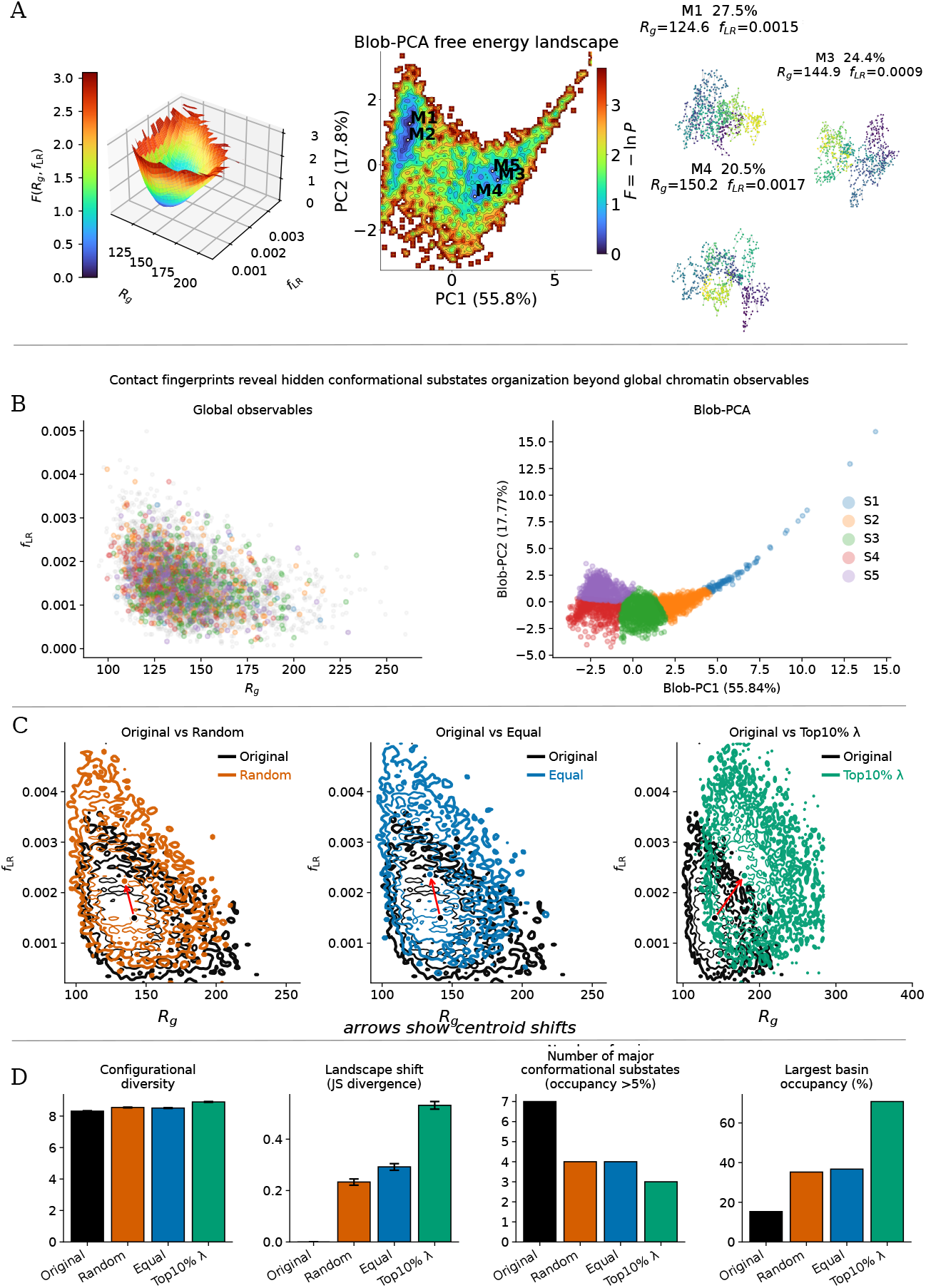
Contact-fingerprint analysis reveals hidden conformational substates and perturbation-dependent reorganisation of the chromatin free-energy landscape. (**A**) Free-energy landscape of the hESC *nanog* locus. Left: three-dimensional surface projected onto global observables, radius of gyration (*R*_*g*_) and long-range contact fraction (*f*_LR_), revealing a single dominant basin. Centre: the same ensemble projected into Blob-PCA contact-fingerprint space, exposing multiple distinct conformational substates (S1-S4) with different occupancies despite overlapping *R*_*g*_ and *f*_LR_ values. Right: representative conformations from substates S1 (27.5%, *R*_*g*_ = 124.6 nm), S3 (24.4%, *R*_*g*_ = 144.9 nm), and S4 (20.5%, *R*_*g*_ = 150.2 nm). **B**) Conformations coloured by Blob-PCA substate identity (S1-S5), projected in global observable space (*R*_*g*_ vs *f*_LR_; left) and contact-fingerprint space (right). Substates that overlap extensively in global observable space are separated in contact-fingerprint space. (**C**) Free-energy landscape contours (*R*_*g*_ vs *f*_LR_) comparing the original MNase-derived ensemble with randomised nucleosome positioning, uniformly spaced nucleosomes, and retention of only the strongest 10% of *λ* interactions. Arrows indicate centroid shifts relative to the original ensemble. (**D**) Quantitative landscape descriptors across all four models: configurational diversity (effective number of occupied states), landscape shift measured by Jensen-Shannon (JS) divergence from the original ensemble, number of major conformational substates (occupancy *>*5%), and largest basin occupancy (fraction of conformations in the most populated substate). Together these metrics distinguish perturbation-induced changes in the location of the free-energy basin from changes in its internal conformational organisation.

This point is made directly by the substate comparison in Fig. 6B. When conformations labelled by their Blob-PCA substate identity (S1-S5) are plotted in *R*_*g*_-*f*_LR_ space, the five substates are completely intermixed and cannot be separated (Fig. 6B, left). The same conformations plotted in Blob-PCA space resolve into non-overlapping clusters (Fig. 6B, right). These substates represent alternative chromatin architectures available to the same genomic locus and therefore constitute a previously unresolved layer of structural variability beyond the ensemble-averaged contact map. Consequently, analyses based solely on compaction or contact-frequency statistics substantially underestimate the structural diversity encoded by the inferred interaction landscape.

To determine how the different components of the *λ* landscape contribute to this free-energy organisation, we compared the original ensemble with three perturbation models: randomised nucleosome positioning, uniformly spaced nucleosomes, and a model retaining only the top 10% of inferred *λ* interactions (Fig. 6C,D). All three perturbations shift the free-energy landscape, but in qualitatively distinct ways that illuminate the separate roles of nucleosome positioning and interaction strength in determining chromatin structural organisation.

The randomised and uniform nucleosome-positioning models both shift the ensemble centroid towards lower *R*_*g*_ and higher *f*_LR_ values relative to the original ensemble (Fig. 6C, left and centre), indicating that disruption of biological nucleosome positioning reduces large-scale chromatin extension while increasing the prevalence of long-range contacts. This is consistent with loss of the locally compact, heterogeneous domain structure generated by realistic nucleosome placement. In contrast, retaining only the top 10% of *λ* interactions shifts the ensemble centroid towards substantially larger *R*_*g*_ and higher *f*_LR_ simultaneously, with the distribution spreading dramatically across a wider conformational space than any nucleosome perturbation condition (Fig. 6C, right). This divergent behaviour confirms that truncating the interaction spectrum distorts the free energy landscape in a qualitatively different manner from nucleosome repositioning. Quantitatively, the Top10% *λ* condition produces by far the largest Jensen-Shannon divergence from the original ensemble (Fig. 6D, landscape shift panel), substantially exceeding both nucleosome perturbation models. All three perturbation conditions reduce the number of major conformational substates relative to the original ensemble (Original ≈ 7; Random, Equal ≈ 4; Top10% ≈ 3; Fig. 6D, third panel). Correspondingly, the largest basin occupancy increases, most severely under Top10% *λ* retention, where the ensemble collapses towards a single dominant substate (≈ 65% occupancy versus ≈ 15% in the original; Fig. 6D, right panel). Together, these results demonstrate that faithful representation of the conformational substate structure requires both biologically realistic nucleosome positioning and the full *λ* interaction spectrum.

The top-10% *λ* truncation model behaves differently, and the contrast with the nucleosome perturbation results is conceptually important (Fig. 6C, right). The centroid shift is modest and the landscape contours retain their overall shape, consistent with Fig. 3, where the strongest 10% of interactions were sufficient to reconstruct the experimental contact map and domain architecture. Here, however, the same truncation fails to reproduce the full conformational diversity of the ensemble. The truncation model shows a striking increase in the largest basin occupancy, defined as the fraction of conformations belonging to the most populated contact-fingerprint substate, and a concomitant reduction in the number of major conformational substates (Fig. 6D). This dissociation between contact map reconstruction and ensemble reconstruction is a key finding with a direct statistical mechanical interpretation: a contact map encodes only the pairwise marginal averages ⟨*h*(*r*_*ij*_) ⟩ for each locus pair, and reproducing these marginals constrains the first moment of each pairwise contact distribution but leaves the joint distribution over all contacts — which determines the conformational ensemble and its free-energy landscape — underdetermined. The same sparse interaction backbone that suffices to recover average contact statistics is therefore insufficient to recover the full distribution of chromatin architectures accessible to the locus. Weak interactions, though dispensable for contact map fidelity, are collectively required to sustain the conformational diversity of the ensemble.

Together, these results reveal a two-tier organisation of structural information within the inferred *λ* landscape. The strongest interactions establish the location and shape of the dominant free-energy basin and are sufficient to recover global chromatin topology and average contact statistics. Weaker interactions encode the internal organisation of that basin, for example, the number and relative occupancy of conformational substates that are invisible to global observables but resolved by contact fingerprint analysis. The *λ* landscape therefore acts not only as a generator of contact frequencies but as a compact representation of the chromatin free-energy landscape itself, simultaneously encoding structural stability, conformational diversity, and the distribution of accessible chromatin architectures at a given locus.

### Targeted perturbation of CTCF/RAD21-associated *λ* interactions reveals a biologically organised architectural backbone

The preceding analyses established that the inferred *λ* landscape encodes a sparse architectural backbone enriched for biologically meaningful chromatin features. A critical question therefore, arises that do the *λ* interactions associated with CTCF/RAD21 binding sites play a structurally privileged role, or do they merely reflect the statistical properties of high-magnitude interactions at any genomic location? To address this, we performed targeted perturbations of the converged *λ* landscape at the K562 *MYC* locus - selectively setting CTCF/RAD21-associated *λ* interactions to zero - and compared their structural consequences against carefully designed magnitude-matched random controls.

CTCF/RAD21-associated *λ* interactions were identified at positions of coincident CTCF and RAD21 enrichment across the locus (Fig. 7A). High-magnitude *λ*_*ij*_ interactions at these positions were selectively set to zero while the remainder of the landscape was left unchanged(Fig. 7B). As a control, an equal number of *λ* interactions were removed from genomic positions lacking CTCF/RAD21 enrichment but matched for both genomic separation and interaction magnitude(Fig. 7B). This matched design is essential. By equating the number and magnitude of removed interactions between conditions, any difference in outcome can be attributed specifically to the biological context of the perturbed interactions, which is their association with CTCF/RAD21 binding, rather than to their statistical properties.

**Fig. 7.**
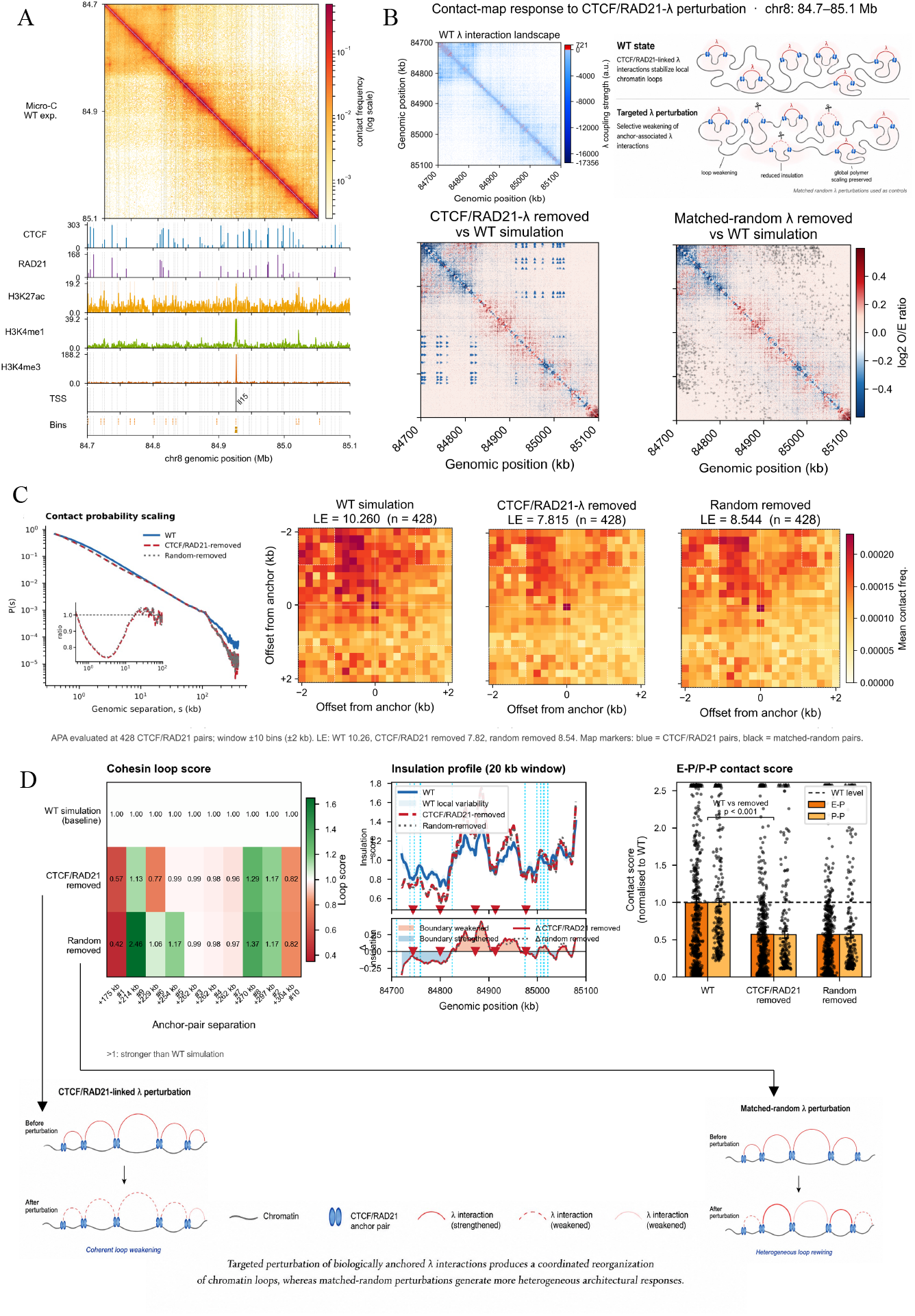
Perturbation of CTCF/RAD21-associated *λ* interactions produces coherent loop weakening while preserving global polymer organisation, in contrast to matched-random perturbations. (**A**) Multi-omic browse rview of the K562 *MYC* locus (chr8: 84.7-85.1 Mb): experimental Micro-C contact map with CTCF, RAD21, H3K27ac, H3K4me1, H3K4me3, and TSS profiles used to define CTCF/RAD21-associated *λ* interactions for targeted perturbation. (**B**) Top left: inferred *λ* interaction landscape for the wild-type (WT) ensemble. Top right: schematic of the perturbation design (targeted CTCF/RAD21-linked *λ* removal versus magnitude-matched random removal at non-CTCF/RAD21 positions). Bottom: log_2_(O/E) contact-ratio maps for CTCF/RAD21-*λ* perturbation (left) and matched-random perturbation (right) relative to WT. (**C**) Left: contact probability scaling *P* (*s*) for WT, CTCF/RAD21-removed, and random-removed ensembles. Centre and right: aggregate peak analysis (APA) centred on 428 CTCF/RAD21 anchor pairs (±2 kb window) for all three conditions. (**D**) Left: cohesin loop scores for individual anchor pairs, normalised to WT, under both perturbation conditions. Centre: insulation profiles (20 kb window) with Δinsulation tracks. Right: enhancer-promoter (E-P) and promoter-promoter (P-P) contact scores normalised to WT. Bottom: schematics of the contrasting architectural responses under targeted and matched-random perturbation.

Contact probability scaling curves *P* (*s*) reveal a distance-dependent pattern of perturbation effects (Fig. 7C, left). At short genomic separations (*s* ≲ 10 kb), all three curves - wild-type (WT), CTCF/RAD21-removed, and random-removed, remain closely superimposed, indicating that local polymer compaction is preserved under both perturbation conditions. At intermediate separations, the inset ratio plot reveals modest but non-zero deviations from the wild-type baseline in both perturbation conditions, with CTCF/RAD21 removal producing a slight reduction in contact probability relative to WT. At large genomic separations (*s* ≳ 100 kb), both perturbation curves fall measurably below the wild-type, indicating a reduction in long-range contact probability that is not captured by the short-range statistics alone. Importantly, these deviations are comparable in magnitude between the CTCF/RAD21-removed and random-removed conditions, suggesting that the reduction in long-range *P* (*s*) reflects a generic consequence of removing a subset of *λ* interactions rather than a specific architectural effect of CTCF/RAD21 loss. The specificity of the CTCF/RAD21 perturbation is therefore not manifest in global *P* (*s*) but is instead concentrated in loop-level and boundary-level reorganisation, as quantified by the APA scores and insulation profiles in Fig. 7C and D.

Against this background of preserved global statistics, targeted removal of CTCF/RAD21-associated *λ* interactions induces a pronounced and spatially organised reorganisation of local chromatin architecture. The log_2_(O/E) contact-ratio map comparing the perturbed and wild-type simulations reveals a structured pattern of contact redistribution concentrated around positions of CTCF/RAD21 association, with coherent domains of contact loss flanked by compensatory contact gain (Fig. 7B, bottom left). This spatial coherence is absent from the matched-random perturbation, which produces a diffuse and heterogeneous contact-ratio map without organised redistribution (Fig. 7B, bottom right). The qualitative difference between these two ratio maps directly demonstrates that CTCF/RAD21-associated *λ* interactions occupy structurally privileged positions in the interaction landscape that cannot be substituted by arbitrary interactions of comparable magnitude at non-CTCF/RAD21 positions.

This conclusion is supported quantitatively by aggregate peak analysis (Fig. 7C, centre and right). Evaluated across 428 CTCF/RAD21 anchor pairs, loop enrichment scores decrease from 10.26 in the wild-type ensemble to 7.82 following targeted perturbation, compared with 8.54 after matched-random perturbation. The greater reduction observed for targeted perturbation - despite identical numbers and magnitudes of removed interactions - establishes that CTCF/RAD21-associated *λ* interactions contribute disproportionately to loop stabilisation relative to interactions at non-CTCF/RAD21 positions.

Analysis of individual loop scores and local insulation further characterises the nature of the architectural response (Fig. 7D). Loop scores across individual anchor pairs reveal a heterogeneous pattern of weakening: some CTCF/RAD21-associated interactions are substantially disrupted while others remain comparatively stable. This heterogeneity suggests that architectural information is not distributed uniformly across all CTCF/RAD21-linked interactions. Instead, a subset of these interactions appears to function as disproportionately important architectural constraints, consistent with the hierarchical organisation of the *λ* backbone established in the truncation analysis of Fig. 3. Insulation profiles show localised boundary weakening near positions of CTCF/RAD21-associated *λ* removal under targeted perturbation, while enhancer-promoter and promoter-promoter contact scores show no significant systematic reduction under either perturbation condition (Fig. 7D, right). The dominant consequence of targeted perturbation is therefore selective disruption of loop architecture and local insulation rather than a broad collapse of chromatin organisation - a distinction that would be obscured if only global contact statistics were examined.

Together, these results provide direct evidence that the inferred *λ* landscape contains a biologically organised architectural backbone in which the spatial context of interactions, not merely their magnitude, determines their structural function. CTCF/RAD21-associated *λ* interactions are not interchangeable with magnitude-matched interactions at arbitrary genomic positions: their selective removal produces coherent loop weakening and localised insulation loss, whereas matched-random removal produces heterogeneous loop rewiring without the same coordinated architectural response. These results complete the central picture emerging from the preceding analyses. The inferred *λ* landscape is not merely a compressed representation of contact frequencies; rather, it encodes biologically organised interaction constraints whose perturbation produces predictable and spatially coherent architectural consequences. Together with the nucleosome-positioning, regulatory-state, and free-energy landscape analyses presented above, this establishes *λ* as a mechanistically interpretable layer of genome organisation beyond contact maps alone.

## Discussion

The central advance of this study is the identification of a nucleosome-resolved interaction landscape *λ* as a physically interpretable layer of chromatin organisation that lies between experimental contact maps and three-dimensional structural ensembles. Contact maps are dominated by polymer-driven distance effects and cannot distinguish enriched from depleted interactions by sign. Structural ensembles are downstream consequences of the interaction field and contain that field only implicitly. *λ* occupies a qualitatively distinct representational role: it encodes the minimal set of pairwise interaction constraints required to explain the observed contact statistics, corrects for genomic distance, concentrates biological signal into a sparse and heavy-tailed backbone, and decomposes into attractive and repulsive components that are separately interpretable in terms of regulatory chromatin state. That a single inference procedure simultaneously achieves distance debiasing, regulatory discrimination, and architectural perturbation prediction - without requiring any epigenomic annotation as input - is the key conceptual advance over prior contact-map analysis methods.

This representational position is conceptually distinct from prior maximum entropy chromatin models. MiChroM and related frameworks treat Lagrange multipliers as energy parameters within a specific interaction-type model and deliver structural ensembles as their primary output [26, 33]. DIMES uses maximum entropy with pairwise distance constraints from imaging data and likewise targets structural ensembles at kilobase-to-chromosome scale [27]. Both are genuinely complementary to the present work: they address the problem of 3D structure reconstruction from different data types and at different resolutions. The distinguishing feature of our framework is that *λ* is the deliverable rather than the conformations it generates. This shift in objective enables the sign decomposition that identifies regulatory state and the sensitivity to nucleosome positioning that contact maps suppress, neither of which is accessible from a structural ensemble alone.

A conceptually significant finding is that nucleosome positioning contributes directly to the interaction landscape in a manner that is substantially invisible to contact maps. The controlled perturbation experiments demonstrate that *λ* responds systematically to nucleosome rearrangements that leave contact maps comparatively unchanged, and that the dominant signal of biologically realistic nucleosome positioning is not a uniform shift in chromatin compaction but an increase in structural heterogeneity - a diversity of local domain sizes and packing states that is absent when positional information is destroyed. This finding connects to emerging experimental evidence that nucleosome positioning encodes 3D genome organisational principles independently of architectural proteins [20, 23, 24], and extends these observations by demonstrating that this positional information is captured in the inferred interaction field rather than in conventional contact statistics. This connection is reinforced by the recent demonstration that native nucleosomes intrinsically encode 3D genome organisation principles independently of architectural proteins [20]. The *λ* framework now provides the quantitative inference machinery to read out that positional encoding from high-resolution contact data. The implication is that contact maps, even at nucleosome resolution, discard a quantifiable amount of structural information that is preserved in *λ*.

The sign decomposition of *λ* into attractive and repulsive components provides a framework for interpreting regulatory chromatin state that is mechanistically grounded in polymer physics rather than empirically derived from epigenomic annotations. The finding that repulsive interactions associate with Polycomb-marked silenced chromatin [40] while attractive interactions concentrate at DNase-accessible, H3K27ac-enriched regulatory regions implies that transcriptional activation and repression are not merely correlated with contact frequency but encoded in the signed interaction field. Regions that make contacts but are governed by repulsive *λ*_*ij*_ values are not held together by specific organisational constraints but by polymer proximity alone - a distinction with direct consequences for predicting the effects of regulatory perturbations such as transcription factor depletion or cohesin loss. The structured CTCF/RAD21 perturbation results provide direct evidence that the architectural backbone inferred by *λ* encodes biologically coordinated spatial organisation: targeted disruption of CTCF/RAD21-associated *λ* interactions selectively redistributes loop enrichment and insulation in ways that magnitude-matched random perturbations cannot reproduce [41, 42], confirming that the spatial context of interactions within the *λ* landscape, not merely their magnitude, determines their structural function.

Several limitations of the present framework should be acknowledged. The inference assumes an effective equilibrium description of chromatin organisation. Chromatin in living cells is driven by active, non-equilibrium processes including transcriptional bursting, loop extrusion dynamics [13], and ATP-dependent chromatin remodelling. However, Micro-C experiments measure population-averaged contact frequencies that represent time-averaged structural statistics. This time-averaging over millions of cells effectively marginalises non-equilibrium single-cell dynamics, producing steady-state contact statistics that are well-approximated by an equilibrium ensemble — an assumption standard across the MaxEnt chromatin modelling literature including MiChroM and DIMES. Under these conditions, equilibrium ensemble inference provides a principled and tractable approach to reconstructing the effective interaction landscape [33, 35, 43]. The inferred *λ*_*ij*_ values should therefore be interpreted as time-averaged effective interaction strengths rather than instantaneous physical forces. A time-resolved extension of the framework, for example using the maximum caliber principle [44], will be required to explicitly capture non-equilibrium chromatin dynamics. The current implementation is computationally limited to locus-scale regions (∼0.2 Mb) at nucleosome resolution. Extension to larger genomic regions will require hierarchical coarse-graining or surrogate inference strategies.

Furthermore, the present analysis spans 12 loci across two human cell lines. While broader than most comparable nucleosome-resolution modelling studies [34, 35], the scope remains limited. Whether the epigenomic correlations and regulatory asymmetries reported here are conserved across cell types, developmental stages, and species will require application of the framework to additional biological contexts. The most consequential limitation concerns nucleosome heterogeneity. The framework uses a single consensus nucleosome map per locus, whereas true nucleosome positions vary across the cell population. Incorporating coherent single-cell nucleosome occupancy data [45] would enable cell-state-specific interaction landscape inference. This would transform *λ* from a population-level descriptor into a tool for dissecting cell-to-cell structural variability - the most direct path towards connecting the inferred interaction landscape to single-cell gene regulatory dynamics.

Despite these limitations, the results establish a clear principle that information beyond conventional contact maps is accessible from Micro-C data when inference is performed at nucleosome resolution with a physically grounded reference model. The *λ* landscape occupies a previously uncharacterised layer of genome organization - above the level of individual contacts, below the level of 3D structural ensembles - that simultaneously encodes interaction sign, structural hierarchy, regulatory identity, and conformational diversity. As single-cell contact data, time-resolved Micro-C, and high-throughput perturbation experiments become available, the maximum entropy inference framework developed here provides a principled and extensible platform for connecting nucleosome architecture, polymer physics, and regulatory epigenomics. More broadly, these results suggest that the latent interaction landscape inferred from contact data may serve as a quantitative bridge between the linear epigenomic sequence of a locus and its three-dimensional regulatory architecture - a connection that has long been sought but has not yet been accessible from contact statistics alone.

## Methods

### Maximum entropy framework

We infer the chromatin interaction landscape from experimental Micro-C contact maps using the maximum entropy (MaxEnt) principle [28, 39]. The objective is to identify the least-biased probability distribution of chromatin conformations that reproduces experimentally observed pairwise contact probabilities 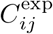 while remaining as close as possible to a physically grounded reference polymer model. The reference distribution *P*_0_(**r**) ∝ exp[−*U*_0_(**r**)] is defined by the polymer energy *U*_0_(**r**) = *U*_bond_ + *U*_bend_ + *U*_ev_, capturing chain connectivity, bending rigidity, and excluded-volume repulsion (Supplementary Methods 1). Maximising the relative entropy of the biased distribution with respect to *P*_0_(**r**) subject to 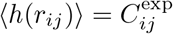 yields the unique MaxEnt distribution (Supplementary Methods 2)

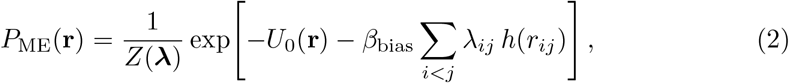

where ***λ*** is the matrix of Lagrange multipliers, *β*_bias_ controls the overall bias strength, and *Z*(***λ***) is the partition function. The contact function *h*(*r*_*ij*_) is a sigmoid with characteristic contact distance *r*_0_ = 30 nm and transition softness *k* = 5 nm. Negative *λ*_*ij*_ corresponds to effective attraction (enriched contacts) and positive to effective repulsion (depleted contacts). To this end, it is noteworthy that the inferred *λ* values are dimensionless Lagrange multipliers and should not be interpreted as physical interaction energies. Their magnitudes reflect the strength of the constraint required to reconcile the experimental contact probability with the reference polymer ensemble. Consequently, large negative values arise for contacts that are substantially enriched relative to polymer expectation, whereas values near zero indicate contacts adequately explained by the reference polymer alone.

### Nucleosome-resolved polymer model

Twelve genomic loci were selected to span a range of transcriptional states: seven from H1-hESC cells (*nanog, cbx8, ppm1g, α*-globin, *hoxa1* as active loci; *hoxb1, hoxc11* as Polycomb-repressed loci) and five from K562 cells (*MYC* as an active locus; *nanog, tal1, lmo2, sox2* as inactive or lowly expressed loci; Supplementary Table 1). Corresponding contact maps are shown in Supplementary Figure 5, demonstrating an excellent match between contact maps generated from MicroC experimental and Maximum entropy inference framework. The recovered interaction landscapes *λ* during Maximum entropy inference framework are presented in Supplementary Figure 6. Each locus is modelled as a heterogeneous coarse-grained copolymer at nucleosome-linker (NL) resolution over a 0.2 Mbp region. Nucleosome positions are derived from MNase-seq data [21, 46] using the DANPOS peak-calling pipeline [37]. Nucleosome core particles are represented as beads of effective diameter ∼10 nm; linker DNA segments are smaller beads of diameter ∼2.5 nm (∼7-8 bp per linker bead). The fine-grained NL chain is grouped into *N*_seg_ contiguous segments (∼200 bp each) matching the experimental Micro-C resolution; contacts are evaluated between segment-pair centres of mass. An initial ensemble of 6,000 random polymer conformations is generated per locus.

### Micro-C contact map preprocessing

Experimental Micro-C contact matrices (Supplementary Figure 5) were balanced using iterative correction and eigenvector decomposition (ICE normalisation) at 200 bp resolution using the cooler software package. Bins with fewer than 10% of the median genomic coverage were masked prior to inference. All contact probabilities were normalised to unit sum within the analysed 0.2 Mb locus window. Robustness of the inferred *λ* landscape to moderate experimental noise was confirmed by re-inference on contact maps perturbed with distance-dependent Gaussian fluctuations (*α* = 0.5), yielding near-unity Spearman correlations with the unperturbed maps at all tested loci (Supplementary Fig. 2-4).

### Iterative inference procedure

The Lagrange multipliers ***λ*** are inferred iteratively through alternating cycles of biased Monte Carlo simulation and multiplier update (Supplementary Fig. 1 for move types). In each cycle, ensemble-averaged contact probabilities 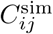 are computed and compared with experimental targets; multipliers are updated by a log-ratio gradient step weighted by genomic distance and contact strength, with L1 sparsity regularisation and hard bounds on |*λ*_*ij*_| (Supplementary Methods 2.3). Periodic relaxation phases allow the ensemble to escape local minima (Supplementary Methods 2.4). Convergence is assessed by stabilisation of 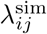 across successive intervals, confirmed by Pearson and Spearman correlations between independent random seeds exceeding 0.99 (Supplementary Figs. 3). Furthermore, the robustness of the Maximum entropy inference framework is tested for two loci by adding gaussian noise to the experimental MicroC map. The recovered contact maps are found to share a high correlation with the experimental MicroC map (CBX8: *r* = 0.924, *ρ* = 0.993; nanog: *r* = 0.881, *ρ* = 0.998), confirming that the inference is not sensitive to moderate experimental perturbations. When tested across all 12 loci, the Pearson correlations at convergence range from 0.77-0.92, with lower values reflecting sparse or weakly structured experimental contact maps in which limited organisational signal above the polymer background correctly drives most multipliers toward zero. Spearman correlations approach 0.99 across all loci confirming faithful recovery of interaction rank ordering. Forward simulations for four genomic loci (Supplementary Fig 8) with fixed converged *λ* using a hamiltonian as give in Eq S12 reproduce these fidelity values closely, and all downstream analyses were performed on loci with Pearson *r* > 0.80. An accurate reproduction of contact probability scaling *P* (*s*) over four decades of genomic separation (Supplementary Figs. 7-8; Supplementary Table 2). Full polymer energy functions, simulation parameters, and all downstream analysis methods (insulation score, blob detection, free-energy landscape, perturbation design, and epigenomic correlation procedures) are provided in Supplementary Methods.

## Data availability

The chromatin contact maps used in this study are derived from previously published datasets. High-resolution Micro-C contact maps for H1-hESC from Krietenstein et al. are available through the 4D Nucleome Data Portal [47] under accession number 4DNES21D8SP8 (https://data.4dnucleome.org/experiment-set-replicates/4DNES21D8SP8/) [9]. Micro-C datasets for K562 were obtained from GEO under accession number GSE206131 [21]. Nucleosome positioning MNase-seq data for hESC were reported by Yazdi et al. (GEO: GSM1194220) [46] and obtained from the NucMap database [48]. The K562 MNase-seq data were obtained from GEO under accession number GSM920557. The mouse celline microc and MNase-seq .The ChIP-seq signal tracks for histone modifications were obtained from the ENCODE Project Consortium.

## Code availability

All custom code for maximum entropy inference, Monte Carlo polymer simulations, and downstream analysis is publicly available at https://github.com/arnabbhattacherjee-lab/maxent-chromatin (DOI to be assigned upon acceptance; a frozen pre-acceptance version has been deposited on Zenodo). Visualisation of chromatin conformations was performed using PyMOL (v2.5.0). Data analysis and plotting used Python (conda environment, key packages: numpy, scipy, matplotlib, cooler) and Gnuplot (v5.4). Schematic figures were prepared using Keynote.

## Acknowledgements

We gratefully acknowledge financial support from DST India (CRG/2023/000636), DBT India (BT/PR46247/BID/7/1015/2023), DBT CoE research grant and JNU ANRF PAIR grant (ANRF/PAIR/2025/000029/PAIR-A). R.M. acknowledges financial support from the Council of Scientific & Industrial Research (CSIR), Govt. of India for Senior Research Fellowship (File No. 09/0263(11815)/2021-EMR-I). The authors acknowledge the generous computational resources provided at the Param Rudra NSM facility hosted by the Inter-University Accelerator Centre (IUAC), New Delhi. All scientific content, analyses, and interpretations were developed solely by the authors.

## Author Contributions

A.B. conceived the study. R.M., K.P.K. and A.B. designed the model and algorithms. R.M. and K.P.K. collected the experimental datasets and performed the simulations. R.M., K.P.K. and A.B. analysed the data and wrote the manuscript.

