## Supplementary File for "A latent interaction landscape inferred from Micro-C encodes chromatin regulatory identity and conformational diversity beyond contact frequency"

**S3 Monte Carlo sampling protocol** **4**

**S4 Locus selection and chromatin state annotation** **4**

**S5 Forward simulation and validation** **6**

**S6 Robustness of inference to experimental noise** **6**

**S7 Observed/expected normalisation and $\lambda$ : a quantitative comparison** **6**

**S8 Insulation score analysis and domain boundary identification** **7**

**S9 Nucleosome-interaction domain (blob) analysis** **7**

### Supplementary Notes

### S1 Nucleosome-resolved polymer model

#### S1.1 Polymer representation

Each genomic locus is modelled as a heterogeneous bead-spring copolymer at nucleosome-linker (NL) resolution spanning ∼0.2 Mb. Nucleosome core particles are represented as beads of effective diameter $\sigma_{\text{nuc}} \approx 10$ nm; each intervening linker segment is a bead of diameter $\sigma_{\text{link}} \approx 2.5$ nm corresponding to ∼7-8 bp of linker DNA. Nucleosome positions are derived from MNase-seq data using the DANPOS peak-calling pipeline, which identifies consensus dyad locations from aligned reads. This representation encodes locus-specific nucleosome spacing and linker-length heterogeneity directly into the reference polymer prior to inference.

For inference and contact-map comparison, the fine-grained NL chain is grouped into $N_{\text{seg}}$ contiguous genomic segments of ∼200 bp, matching the experimental Micro-C resolution. Contacts are evaluated between segment centres of mass, enabling direct comparison with experimental contact probabilities.

The total energy of a conformation $\mathbf{r}$ is

$$U_{\text{total}}(\mathbf{r}) = U_{\text{bond}}(\mathbf{r}) + U_{\text{bend}}(\mathbf{r}) + U_{\text{ev}}(\mathbf{r}) + U_{\text{bias}}(\mathbf{r}), \tag{S1}$$

where the first three terms constitute the reference polymer energy $U_0(\mathbf{r})$ and the last term encodes the MaxEnt bias (Section S2.2).

**Bond potential.** Adjacent beads are connected by harmonic springs:

$$U_{\text{bond}} = \sum_{i=1}^{N-1} \frac{k_{\text{bond}}}{2} \left(r_{i,i+1} - r_0\right)^2, \tag{S2}$$

where $r_0$ is the equilibrium bond length and $k_{\text{bond}}$ is the spring constant. For nucleosome-nucleosome bonds $r_0 = 10$ nm; for linker-to-nucleosome bonds $r_0$ is set proportional to the linker length.

**Bending potential.** Chain stiffness is introduced through a cosine bending energy:

$$U_{\text{bend}} = k_{\text{bend}} \sum_{i=2}^{N-1} (1 - \cos\theta_i), \tag{S3}$$

where $\theta_i$ is the angle between consecutive bond vectors. The persistence length of the chain is set to be consistent with the experimentally measured chromatin fibre stiffness of ∼50 nm.

**Excluded-volume interaction.** Steric repulsion between non-bonded beads is modelled by the purely repulsive (truncated and shifted) Lennard-Jones potential:

$$U_{\text{ev}}(r_{ij}) = \begin{cases} \epsilon \left[ \left(\frac{\sigma_{ij}}{r_{ij}}\right)^{12} - \left(\frac{\sigma_{ij}}{r_{ij}}\right)^{6} - U(r_c) \right], & r_{ij} < r_c, \\ 0, & r_{ij} \geq r_c, \end{cases} \tag{S4}$$

where $\sigma_{ij} = (\sigma_i + \sigma_j)/2$ is the mixed diameter, $\epsilon$ sets the energy scale, and the cutoff $r_c = 2^{1/6}\sigma_{ij}$ ensures that only the repulsive wall is active. This term prevents bead overlap while imposing no spurious long-range attractions.

#### S1.2 Contact function

The contact probability between genomic segments $i$ and $j$ is encoded by the sigmoid function

$$h(r_{ij}) = \frac{1}{1 + \exp[(r_{ij} - r_0^c)/k_s]}, \tag{S5}$$

where $r_0^c = 30$ nm is the characteristic contact distance and $k_s = 5$ nm controls the transition softness. This continuous form avoids hard-cutoff discontinuities and is well suited to gradient-based optimisation. The ensemble-averaged simulated contact probability is

$$C_{ij}^{\mathrm{sim}} = \frac{1}{T_{\mathrm{snap}}} \sum_{t=1}^{T_{\mathrm{snap}}} h\Big(r_{ij}^{(t)}\Big), \tag{S6}$$

where $T_{\mathrm{snap}}$ is the number of sampled conformations.

### S2 Maximum entropy inference framework

#### S2.1 Theoretical basis

We seek the probability distribution $P(\mathbf{r})$ over chromatin conformations that (i) reproduces the experimental contact probabilities $C_{ij}^{\mathrm{exp}}$ and (ii) is maximally non-committal with respect to all other information, i.e., is closest to the reference polymer distribution $P_0(\mathbf{r}) \propto \exp[-U_0(\mathbf{r})]$ in the sense of minimum relative entropy (Kullback-Leibler divergence).

By the method of Lagrange multipliers, the unique solution is:

$$P_{\mathrm{ME}}(\mathbf{r}) = \frac{1}{Z(\boldsymbol{\lambda})} \exp\left[-U_0(\mathbf{r}) - \beta_{\mathrm{bias}} \sum_{i<j} \lambda_{ij}\, h(r_{ij})\right], \tag{S7}$$

where $\boldsymbol{\lambda}$ is the matrix of Lagrange multipliers, $\beta_{\mathrm{bias}}$ controls the overall strength of the bias, and $Z(\boldsymbol{\lambda})$ is the partition function. The MaxEnt constraints $\langle h(r_{ij})\rangle_\lambda = C_{ij}^{\mathrm{exp}}$ uniquely determine $\boldsymbol{\lambda}$.

Negative $\lambda_{ij}$ values correspond to effective attraction between segments $i$ and $j$ (contacts enriched relative to the reference polymer), while positive values correspond to effective repulsion (contacts depleted below the reference expectation). A near-zero $\lambda_{ij}$ indicates that the reference polymer alone accounts for the observed contact probability.

#### S2.2 MaxEnt bias potential

The bias term in Eq. (S1) is

$$U_{\mathrm{bias}}(\mathbf{r}) = \beta_{\mathrm{bias}} \sum_{i<j} \lambda_{ij}\, h(r_{ij}). \tag{S8}$$

A repulsive penalty is additionally applied when a segment pair with no experimental contact ($C_{ij}^{\mathrm{exp}} \approx 0$) approaches closer than a threshold:

$$U_{\mathrm{rep}}(r_{ij}) = \frac{5\, C_{ij}^{\mathrm{sim}}}{(r_{ij} - 1.5\, r_c)^2}, \qquad r_{ij} < 1.5\, r_c, \tag{S9}$$

which suppresses spurious contacts not supported by the experimental data.

#### S2.3 Iterative update of Lagrange multipliers

The multipliers $\lambda_{ij}$ are inferred iteratively. After each bias-update interval, the ensemble-averaged contact map $C_{ij}^{\mathrm{sim}}$ is recomputed and the multipliers are updated by a scale-aware, distance-weighted logarithmic gradient step:

$$\lambda_{ij}^{(t+1)} = \lambda_{ij}^{(t)} + \eta\, g_{ij}\, s_{ij} \ln\left(\frac{C_{ij}^{\mathrm{sim}} + \epsilon}{C_{ij}^{\mathrm{exp}} + \epsilon}\right) - \eta\, \lambda_{L_1} \operatorname{sign}(\lambda_{ij}), \tag{S10}$$

where:

- $\eta$ is the learning rate;
- $g_{ij}$ is a genomic-distance-dependent weight that down-weights noisy long-range contacts;
- $s_{ij} = 1 + C_{ij}^{\mathrm{exp}}$ is a strength factor that amplifies updates for high-confidence contacts;
- $\epsilon \ll 1$ ensures numerical stability near zero;

- the $L_1$ regularisation term ($\lambda_{L_1}$) promotes sparsity by driving weak, unsupported interactions toward zero.

The logarithmic form ensures that updates scale with relative rather than absolute deviations, improving convergence across the many orders of magnitude spanned by contact probabilities. Multipliers are hard-bounded at $|\lambda_{ij}| \leq \lambda_{\max}$, and entries with $|\lambda_{ij}| < 10^{-4}$ are thresholded to zero to eliminate negligible interactions.

##### S2.4 Bias-relax optimisation schedule

To avoid entrapment in metastable states and improve convergence, the inference proceeds through three sequential phases applied cyclically (100 cycles per locus):

**Phase 0 - Bias ramping.** The bias strength $\beta_{\text{bias}}$ is increased multiplicatively ($\beta_{\text{bias}} \leftarrow 1.005\,\beta_{\text{bias}}$) until the target maximum $\beta_{\max}$ is reached, allowing the ensemble to adapt smoothly to the imposed contact constraints.

**Phase 1 - Relaxation.** The bias is temporarily reduced ($\beta_{\text{bias}} \sim 10^{-3}$) and the simulation temperature is raised ($T = 2.0$), promoting broad exploration of conformational space and preventing overfitting to the current $\boldsymbol{\lambda}$ estimate. The multipliers are weakly damped during this phase.

**Phase 2 - Re-biasing and refinement.** The bias is reintroduced and the multipliers are updated using Eq. (S10). Transitions between phases are governed by predefined MC-step counts and convergence criteria (see below).

##### S2.5 Convergence assessment

Convergence is assessed by (i) stabilisation of $C_{ij}^{\text{sim}}$ across successive optimisation intervals and (ii) Pearson and Spearman correlations between simulated contact maps computed from independent random seeds exceeding 0.99. Supplementary Fig. 2 shows representative convergence traces for all 12 analysed loci.

#### S3 Monte Carlo sampling protocol

Structural ensembles consistent with $P_{\text{ME}}(\mathbf{r})$ are generated by parallel Metropolis Monte Carlo (MC) at inverse temperature $\beta = 1$. An initial ensemble of 6,000 independent polymer conformations is evolved simultaneously. Trial moves are drawn from three complementary move types (Supplementary Fig. 1):

**Pivot move.** A randomly selected bead acts as a pivot; a contiguous polymer segment is rotated about a random axis by a uniformly distributed angle. This produces large-scale rearrangements and ensures efficient exploration of global conformational space.

**Crankshaft rotation.** A short sub-chain is rotated about the axis connecting its two endpoints, producing local conformational changes while preserving bond geometry.

**Gaussian displacement.** A short segment is displaced by a small Gaussian-distributed random vector, enabling fine-scale adjustments and local energy minimisation.

A trial move generating candidate conformation $\mathbf{r}'$ is accepted with probability

$$P_{\text{acc}} = \min(1, \exp[-\Delta U])\,, \quad \Delta U = U(\mathbf{r}') - U(\mathbf{r}), \tag{S11}$$

which satisfies detailed balance and guarantees convergence to the target distribution. Sampling is parallelised across independent snapshots using OpenMP; global observables (ensemble-averaged contact maps) are synchronised at regular intervals.

#### S4 Locus selection and chromatin state annotation

Twelve genomic loci ($\sim$0.2 Mb each) were selected to span a range of transcriptional states in two human cell lines:

- **H1-hESC (7 loci):** nanog, cbx8, ppm1g, $\alpha$-globin, hoxa1 (active); hoxb1, hoxc11 (inactive/Polycomb-repressed).

- **K562 (5 loci):** myc (active); nanog, sox2, lmo2, tal1 (inactive or lowly expressed in this lineage).

Transcriptional activity was defined by ChromHMM annotation (15-state model) (Supplementary Table S1).

**Table S1: Genomic loci analysed, their chromosomal coordinates, cell line, and transcriptional state.** Coordinates refer to the GRCh38/hg38 assembly. ChromHMM states: A, active transcription; PC, Polycomb-repressed; L, low/bivalent.

| Locus | Cell line | Coordinates (hg38) | ChromHMM | State |
|---|---|---|---|---|
| ppm1g | hESC | chr2:27.3–27.5 Mb | A | Active |
| $\alpha$-globin | hESC | chr16:0.1–0.3 Mb | A | Active |
| cbx8 | hESC | chr17:79.7–79.9 Mb | A | Active |
| nanog | hESC | chr12:7.7–7.9 Mb | A | Active |
| hoxa1 | hESC | chr7:27.0–27.2 Mb | PC | Inactive |
| hoxb1 | hESC | chr17:48.4–48.6 Mb | PC | Inactive |
| hoxc11 | hESC | chr12:53.9–54.1 Mb | PC | Inactive |
| lmo2 | K562 | chr11:33.8–34.0 Mb | L | Low |
| myc | K562 | chr8:127.6–127.8 Mb | A | Active |
| nanog | K562 | chr12:7.7–7.9 Mb | L | Inactive |
| sox2 | K562 | chr3:181.6–181.8 Mb | L | Low |
| tal1 | K562 | chr1:47.1–47.3 Mb | L | Low |

Pearson correlations at MaxEnt convergence range from 0.77 to 0.92 across the 12 analysed loci. The lower values occur exclusively at loci with sparse or weakly structured experimental Micro-C contact maps, where limited organisational signal above the polymer background results in near-zero inferred multipliers for most locus pairs. In these cases, the observed contact statistics are largely accounted for by the reference polymer model, indicating that only a small subset of interactions requires additional effective constraints. This behaviour is consistent with the maximum entropy formalism, which introduces non-zero interactions only when required to reproduce deviations from the polymer baseline. Despite variation in Pearson correlation, Spearman correlations approach 0.99 across all loci, demonstrating that the relative ordering of interactions is recovered robustly regardless of contact-map sparsity. Forward simulations performed with the converged $\lambda$ landscape held fixed reproduce the MaxEnt Pearson correlations closely, confirming that the fixed-$\lambda$ evaluation introduces no appreciable loss in reconstruction fidelity.

**Table S2: MAXENT-simulation fidelity across all 12 analysed loci.** Pearson ($r$) and Spearman ($\rho$) correlations between forward-simulated and experimental Micro-C contact maps. hESC stands for human embryonic stem cells.

| Locus | Cell line | $r$ | $\rho$ |
|---|---|---|---|
| nanog | hESC | 0.92 | 0.99 |
| cbx8 | hESC | 0.91 | 0.99 |
| ppm1g | hESC | 0.89 | 0.99 |
| $\alpha$-globin | hESC | 0.88 | 0.99 |
| hoxa1 | hESC | 0.85 | 0.99 |
| hoxb1 | hESC | 0.83 | 0.99 |
| hoxc11 | hESC | 0.81 | 0.99 |
| myc | K562 | 0.90 | 0.99 |
| nanog | K562 | 0.86 | 0.99 |
| tal1 | K562 | 0.84 | 0.99 |
| lmo2 | K562 | 0.80 | 0.99 |
| sox2 | K562 | 0.77 | 0.99 |

### S5 Forward simulation and validation

To validate the inferred $\boldsymbol{\lambda}$, we perform forward Monte Carlo simulations with the converged multipliers held fixed. No parameter updates are applied; the system evolves under

$$H(\mathbf{r}) = U_0(\mathbf{r}) + \beta_{\text{bias}} \sum_{i<j} \lambda_{ij}\, h(r_{ij}), \tag{S12}$$

and the ensemble-averaged contact map

$$\langle C_{ij} \rangle = \frac{1}{T_{\text{snap}}} \sum_{m=1}^{T_{\text{snap}}} h\left(r_{ij}^{(m)}\right) \tag{S13}$$

is compared directly to $C_{ij}^{\text{exp}}$.

Across all 12 loci, Pearson correlations between forward-simulated and experimental contact maps range from 0.77 to 0.92; Spearman correlations approach 0.99 (Supplementary Table S2). The forward simulations also accurately reproduce the contact probability scaling $P(s)$ over four decades of genomic separation (Supplementary Fig. 7), confirming that the inferred landscape captures both local and global structural features without overfitting.

### S6 Robustness of inference to experimental noise

To assess robustness of the MaxEnt framework against experimental noise, we generated perturbed contact maps by adding distance-dependent Gaussian fluctuations to the experimental Micro-C matrices:

$$C_{ij}^{\text{noise}} = \max(0,\ C_{ij} + \mathcal{N}(0, \alpha\sigma_d))\,, \tag{S14}$$

where $\sigma_d$ is the empirical standard deviation of all contact values at genomic separation $d = |i - j|$ and $\alpha = 0.5$ introduces moderate noise while preserving distance-dependent statistics. Negative values are truncated to zero to maintain physical contact frequencies.

Perturbed maps were used as input to the full inference pipeline. At the ppm1g locus, the Pearson/Spearman correlations between the noisy-input simulated map and the original experimental map were 0.924/0.993; at the nanog locus, 0.881/0.998. Near-unity Spearman correlations across both loci confirm that the inferred interaction landscape is robust to moderate experimental perturbations and does not overfit to noise in the input contact data (Supplementary Fig. 4).

### S7 Observed/expected normalisation and $\lambda$ : a quantitative comparison

A key question is whether the inferred $\lambda$ landscape constitutes genuinely new information or merely reparameterises standard observed/expected (O/E) contact normalisation. We address this quantitatively by comparing $|\lambda_{ij}|$ against raw and O/E-normalised contacts across all segment pairs.

The Pearson correlation between $|\lambda_{ij}|$ and raw contact frequency is $r = 0.262$, rising to $r = 0.539$ upon O/E normalisation, confirming that $\lambda$ captures the distance-corrected interaction structure more faithfully than raw frequency. However, three properties of $\lambda$ are inaccessible to O/E normalisation:

1. **Sign.** O/E maps are non-negative by construction and cannot distinguish enriched ($\lambda^- < 0$) from depleted ($\lambda^+ > 0$) interactions. The sign decomposition of $\lambda$ identifies regulatory state independently of epigenomic annotation (main text, Section 1.5).

2. **Sparsity.** The $\lambda$ matrix carries ~90% near-zero entries, concentrating organisational signal into a sparse high-confidence interaction backbone; O/E maps are dense.

3. **Spatial correlations.** Distance-preserving shuffling of contact probabilities destroys the coherent domain-like spatial patterns in $\lambda$ while leaving O/E statistics unchanged (Supplementary Fig. 10).

Together these comparisons demonstrate that $\lambda$ is a qualitatively distinct representation grounded in the polymer physics of the reference ensemble, not a reparameterisation of O/E-normalised contacts.

### S8 Insulation score analysis and domain boundary identification

Domain boundaries were identified using the insulation score method [?]. For a given contact map, the insulation score at genomic position $k$ is computed as

$$\mathrm{IS}_k = \frac{1}{2d} \sum_{i=k-d+1}^{k} \sum_{j=k+1}^{k+d} x_{ij}, \qquad d < k < N - d, \tag{S15}$$

where $x_{ij}$ is the contact frequency between segments $i$ and $j$, $d$ is the half-window size, and $N$ is the total number of segments. A position is classified as a boundary if its insulation score constitutes a local minimum relative to a comparison window of $K$ bins on each side. Parameters $d$ and $K$ were chosen based on visual inspection and cross-validated against experimental TAD calls from published studies.

Two complementary metrics are reported in main text Fig. 3C:

**Overlap fraction.** The proportion of experimentally detected boundaries that are positionally recovered within a tolerance window of $\delta$ bp.

**Pearson correlation of insulation scores.** The Pearson correlation between experimental and simulated insulation score values at matched boundary positions, quantifying the fidelity of insulation depth reproduction independently of positional tolerance.

The combined use of these metrics guards against artificially inflated overlap fractions that arise when sparse $\lambda$ truncations produce broad, shallow insulation minima that claim multiple experimental boundaries at low positional stringency.

**Table S3:** Correlation between MNase-Seq and alternative nucleosome positioning models (equi-spaced and random) for both contact maps and $\lambda$ maps across loci and cell lines. Values are Pearson and Spearman correlation coefficients (rounded to two decimal places).

| Map | Locus | Cell line | Comparison | Pearson | Spearman |
|---|---|---|---|---|---|
| Contact | nanog | hESC | MNase vs Equi-Spaced | 0.99 | 0.99 |
| Contact | nanog | hESC | MNase vs Random | 0.99 | 0.99 |
| Contact | hoxc11 | hESC | MNase vs Equi-Spaced | 0.99 | 0.97 |
| Contact | hoxc11 | hESC | MNase vs Random | 0.99 | 0.97 |
| Contact | nanog | K562 | MNase vs Equi-Spaced | 0.99 | 0.99 |
| Contact | nanog | K562 | MNase vs Random | 0.99 | 0.99 |
| Contact | myc | K562 | MNase vs Equi-Spaced | 0.99 | 0.99 |
| Contact | myc | K562 | MNase vs Random | 0.99 | 0.99 |
| $\lambda$ | nanog | hESC | MNase vs Equi-Spaced | 0.99 | 0.92 |
| $\lambda$ | nanog | hESC | MNase vs Random | 0.99 | 0.92 |
| $\lambda$ | hoxc11 | hESC | MNase vs Equi-Spaced | 0.99 | 0.96 |
| $\lambda$ | hoxc11 | hESC | MNase vs Random | 0.99 | 0.96 |
| $\lambda$ | nanog | K562 | MNase vs Equi-Spaced | 0.99 | 0.93 |
| $\lambda$ | nanog | K562 | MNase vs Random | 0.99 | 0.92 |
| $\lambda$ | myc | K562 | MNase vs Equi-Spaced | 0.99 | 0.94 |
| $\lambda$ | myc | K562 | MNase vs Random | 0.99 | 0.93 |

### S9 Nucleosome-interaction domain (blob) analysis

#### S9.1 Blob detection algorithm

Chromatin conformations were partitioned into spatially compact nucleosome interaction domains - hereafter referred to as "blobs" - using the Density-Based Spatial Clustering of Applications with Noise (DBSCAN) algorithm [?], applied directly to the three-dimensional Cartesian coordinates of nucleosome bead centres extracted from each simulation snapshot [?]. DBSCAN groups points into clusters on the basis of two parameters: a neighbourhood radius $\varepsilon$ (nm) and a minimum cluster population $n_{\mathrm{min}}$. A nucleosome bead is assigned to a cluster if at least $n_{\mathrm{min}}$ beads lie within Euclidean distance $\varepsilon$ of it; beads that satisfy neither criterion are labelled as noise and excluded from downstream analysis. DBSCAN is particularly suited to chromatin blob detection because it makes no assumptions about cluster shape or number, naturally handles the irregular, non-convex geometries of nucleosome aggregates, and distinguishes genuine dense-core clusters from low-density peripheral beads.

Parameters were set to $\varepsilon = 24\,\text{nm}$ and $n_{\text{min}} = 4$ nucleosomes, consistent with the characteristic nearest-neighbour distance in nucleosome-linker polymer ensembles and the minimum size required to define a geometrically resolvable three-dimensional domain. Blob detection was applied to every fifth snapshot of the ensemble (stride = 5) to balance computational cost and statistical coverage.

For each detected blob in each snapshot, the following quantities were computed:

- **Blob size** (bp): the total genomic extent of bead positions assigned to the cluster, computed as the sum of nucleosome core lengths (141 bp per nucleosome) and intervening linker lengths (7.35 bp per linker bead), accounting for the discrete bead-to-genomic position mapping.
- **Radius of gyration** $R_g$ (nm): the root-mean-square distance of beads in the cluster from their centroid.
- **Packing density** (bp/nm$^3$): the blob size divided by the convex-hull volume of the cluster.
- **Eccentricity**: computed from the eigenvalues of the gyration tensor of convex-hull vertices as $e = \sqrt{\max(0, 1 - (I_2 + I_3)/(2\,I_1))}$, where $I_1 \geq I_2 \geq I_3$ are the principal moments ordered by decreasing magnitude. Values near unity indicate strongly elongated, ellipsoidal morphology; values near zero indicate near-spherical blobs.
- **Surface area** ($\mu$m$^2$) and **volume** (nm$^3$): estimated from the convex hull of the nucleosome coordinates comprising the blob, using the Quickhull algorithm as implemented in `scipy.spatial.ConvexHull`. Blobs with fewer than four non-coplanar beads were excluded from geometric analysis.

Blob detection was performed independently for all four nucleosome-positioning models analysed (MNase-seq, randomised, uniformly spaced, and top-2% $\lambda$ truncation), enabling direct cross-configuration comparison of blob geometry and size distributions (Supplementary Fig. S12).

#### S9.2 Blob-PCA free-energy landscape analysis

To expose conformational substates of the chromatin ensemble that are invisible to global structural observables such as radius of gyration ($R_g$) and long-range contact fraction ($f_{\text{LR}}$), we developed a Blob-PCA contact-fingerprint approach that characterises the free-energy landscape at the level of individual nucleosome blob geometry rather than whole-locus shape.

**Feature construction.** For each DBSCAN-detected blob, a nine-dimensional feature vector was constructed from the following geometric descriptors:

1. $\log(1 + S_{\text{bp}})$, where $S_{\text{bp}}$ is the blob size in base pairs;
2. eccentricity $e$;
3. minor-to-major axis ratio $c/a$;
4. minor-to-intermediate axis ratio $c/b$;
5. $\log(1 + A_{\text{hull}})$, where $A_{\text{hull}}$ is the convex-hull surface area;
6. $\log(1 + V_{\text{hull}})$, where $V_{\text{hull}}$ is the convex-hull volume;
7. blob radius of gyration $R_g$ (nm);
8. $\log(1 + \rho_{\text{NUC}})$, where $\rho_{\text{NUC}}$ is the nucleosome number density (nucleosomes/nm$^3$);
9. $\log(1 + \rho_{\text{bp}})$, where $\rho_{\text{bp}}$ is the genomic packing density (bp/nm$^3$).

Logarithmic transforms were applied to size, area, volume, and density features to compress their heavy-tailed distributions and normalise their dynamic range prior to dimensionality reduction. Blobs for which any feature was non-finite (arising from degenerate convex hulls with fewer than four non-coplanar beads) were excluded.

**Dimensionality reduction and landscape construction.** Feature vectors from all valid blobs across all four configurations (Original MNase, Randomised, Equal-spaced, Top-2% $\lambda$) were pooled and jointly standardised to zero mean and unit variance using `StandardScaler`. A shared PCA basis was then fitted on this pooled, standardised matrix using two principal components (`sklearn.decomposition.PCA`, `random_state=42`). Fitting on the pooled data ensures that all configurations are embedded in a common coordinate system, making cross-configuration comparisons directly interpretable.

Each configuration's blobs were then projected onto this shared PC basis, yielding per-blob coordinates $(z_1, z_2)$ in Blob-PCA space. A two-dimensional probability density $P(z_1, z_2)$ was estimated by histogramming these projections onto a $120 \times 120$ grid (spanning the 0.25th–99.75th percentile range along PC1 and the 0.5th–99.5th percentile range along PC2, with 10% and 6% padding, respectively), followed by Gaussian smoothing with $\sigma = 1.0$ bin. The dimensionless free-energy surface was computed as

$$F(z_1, z_2) = -\ln P(z_1, z_2), \tag{S16}$$

shifted so that the global minimum within the occupied region (defined as $P > 0.015 \cdot P_{\max}$) equals zero.

**Conformational substate identification and occupancy.** Free-energy minima were identified as local minima of the Gaussian-smoothed surface $\tilde{F}$ within the occupied region, using a minimum filter of footprint $7 \times 7$ bins and an energy cutoff $\tilde{F} < 6.0\, k_{\mathrm{B}}T$. Only minima located at least two bins from the boundary of the grid were retained. Each occupied pixel was assigned to its nearest free-energy minimum (by Euclidean distance in grid coordinates), and the basin occupancy of each substate was computed as the fraction of total probability mass assigned to that basin. Substates with occupancy > 5% were classified as major conformational substates. The largest basin occupancy and the number of major substates were used as quantitative descriptors of conformational diversity across perturbation conditions (Fig. 6D). The Jensen–Shannon divergence between the Original and each perturbed configuration was computed from the two-dimensional probability distributions and used as a scalar measure of landscape shift (Fig. 6D, second panel). All Jensen–Shannon divergences were estimated with $N_{\mathrm{boot}} = 300$ bootstrap replicates (sampling with replacement) to provide 95% confidence intervals.

#### S9.3 Comparison with experimental blob geometry

Simulated blob geometries were compared with those measured by Barth et al. using live super-resolution chromatin imaging [?]. Experimental blob areas follow a log-normal distribution with a mean of $(3.3 \pm 2.8) \times 10^{-3}\ \mu\mathrm{m}^2$, and the eccentricity distribution peaks near ∼0.9, indicating predominantly elongated structures. Our simulations yield a median blob area of $\sim 5 \times 10^{-3}\ \mu\mathrm{m}^2$ (somewhat broader distribution with a pronounced tail) and a median eccentricity of ∼0.9, consistent with the experimental observations (Supplementary Table S4).

**Table S4: Blob geometry across all analysed genomic loci.** Median eccentricity and projected surface area of chromatin blobs for each locus; the final row gives the overall median across all loci. (hESC-human embryonic stem cells; K562-chronic myelogenous leukaemia cells.)

| **Locus** | **Cell line** | **Eccentricity (median)** | **Area (median, $\mu$m$^2$)** |
|---|---|---|---|
| nanog | hESC | 0.913 | 0.00318 |
| ppm1g | hESC | 0.892 | 0.01145 |
| $\alpha$-globin | hESC | 0.899 | 0.00605 |
| cbx8 | hESC | 0.892 | 0.00893 |
| hoxa1 | hESC | 0.905 | 0.00519 |
| hoxb1 | hESC | 0.890 | 0.01248 |
| hoxc11 | hESC | 0.896 | 0.00867 |
| myc | K562 | 0.903 | 0.00686 |
| nanog | K562 | 0.901 | 0.00490 |
| tal1 | K562 | 0.920 | 0.00329 |
| lmo2 | K562 | 0.902 | 0.00674 |
| sox2 | K562 | 0.882 | 0.00712 |
| **Overall** | - | **0.906** | **0.00496** |

### S10 Epigenomic correlation analysis

#### S10.1 Chromatin mark enrichment at high-$|\lambda|$ bins

To test whether $\lambda$ carries regulatory information beyond contact frequency, we compared chromatin mark signal enrichment at high-$|\lambda|$ genomic bins against a contact-frequency-matched baseline. For each ChIP-seq/ATAC-seq track, the following procedure was applied:

1. Genomic bins were ranked by $|\lambda_{ij}|$ summed over all $j$ (the "$\lambda$-weight" of bin $i$).
2. A set of high-$|\lambda|$ bins was defined as the top 20% of bins by $\lambda$-weight.
3. A contact-frequency-matched baseline was constructed by selecting, for each high-$|\lambda|$ bin, a set of bins with matched total contact frequency ($\pm 10\%$) drawn uniformly at random (without replacement) from all remaining bins.
4. Enrichment $\Delta$signal was defined as the mean epigenomic signal at high-$|\lambda|$ bins minus the mean signal at the matched baseline.

This design isolates the regulatory contribution of $\lambda$ from the trivial dependency on contact strength. Separate enrichments were computed for the attractive ($\lambda^-$) and repulsive ($\lambda^+$) components.

#### S10.2 Spearman correlation of $\lambda$ sign with chromatin marks

Spearman correlations were computed between the genomic profiles of $\lambda^-$ or $\lambda^+$ (projected along the linear genome by summing over all interaction partners) and normalised ChIP-seq/ATAC-seq signal tracks for the following epigenomic features: DNase-I hypersensitivity, H3K27ac, H3K4me1, H3K4me3, EP300, POLR2A (active/enhancer marks); CTCF, RAD21 (architectural marks); H3K36me3, H3K27me3 (repressive/elongation marks). Correlation $p$-values were corrected for multiple comparisons using the Benjamini-Hochberg procedure.

#### S10.3 $\lambda$ advantage over contact frequency

Per-mark predictive advantage of $|\lambda|$ over the best-performing contact-based predictor (raw or O/E) was quantified as:

$$\Delta\rho_{\text{mark}} = \rho(|\lambda|, \text{mark}) - \max\Big[\rho(C^{\text{raw}}, \text{mark})\,,\ \rho\Big(C^{\text{O/E}}, \text{mark}\Big)\Big]\,. \tag{S17}$$

Positive values indicate that $\lambda$ outperforms contact-based predictors for a given mark.

### S11 CTCF-linked perturbation analysis

#### S11.1 Perturbation design

To test whether the inferred $\lambda$ landscape encodes a biologically organised architectural backbone beyond generic polymer statistics, we performed two classes of perturbation on the converged K562 myc $\lambda$ matrix:

**CTCF-linked perturbation.** Convergent CTCF binding sites were identified within the myc locus using CTCF ChIP-seq peaks (ENCODE; K562). For each pair of convergently oriented CTCF sites, the high-magnitude $\lambda_{ij}$ interactions centred on those sites (within $\pm 2$ bins of the CTCF peak positions) were identified. The selected interactions were perturbed by setting $\lambda_{ij} \leftarrow 0$ (removal of the interaction constraint), and the forward simulation was re-run with the modified landscape.

**Randomised matched perturbation.** An equal number of $\lambda_{ij}$ interactions were perturbed at positions selected uniformly at random, subject to the constraints that (i) the perturbed interactions have the same genomic-distance distribution and (ii) the same $|\lambda|$ distribution as those perturbed in the CTCF-linked class. This matched-random control ensures that any downstream structural differences arise from the spatial organisation of the perturbed interactions, not their statistical properties.

#### S11.2 Variance-difference analysis

For each perturbation ensemble, the positional variance $\text{Var}(\mathbf{r}_i)$ was computed for every bead $i$. Variance-difference maps $\Delta V_{ij} = \text{Var}_{\text{perturbed}} - \text{Var}_{\text{WT}}$ were then constructed; positive (red) regions indicate enhanced positional fluctuations and negative (blue) regions indicate suppressed fluctuations relative to the wild-type ensemble.

##### S11.3 Principal component analysis and Wasserstein distances

The WT conformational ensemble was projected onto its principal components (PCs) using the covariance matrix of bead coordinates. CTCF-linked and randomised ensembles were then projected onto the same PC basis. Wasserstein distances between WT and perturbed distributions along each PC were computed as the $L_1$ distance between the corresponding empirical cumulative distribution functions.

#### Supplementary Figures

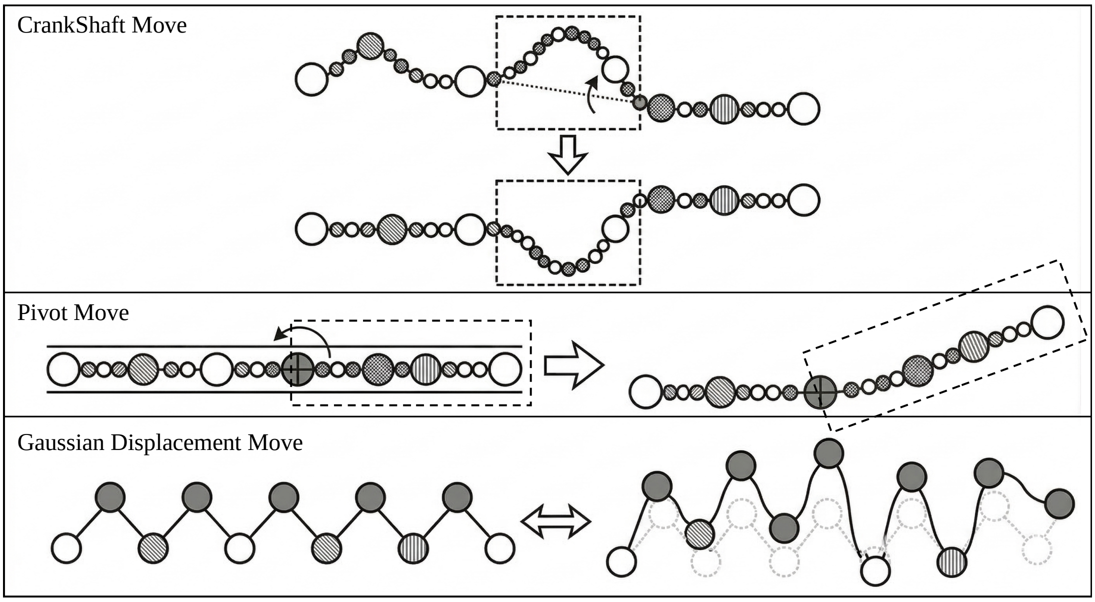

**Figure 1: Monte Carlo move sets used in polymer conformational sampling.** Three complementary move types ensure efficient exploration across multiple length scales. **Top:** Crankshaft rotation - a short sub-chain segment is rotated about the axis connecting its two endpoint beads. **Middle:** Pivot move - a contiguous polymer segment is rotated about a randomly chosen bead by a random angle, enabling large-scale rearrangements. **Bottom:** Gaussian displacement - a local segment is displaced by a small Gaussian-distributed random vector, enabling fine-scale adjustments.

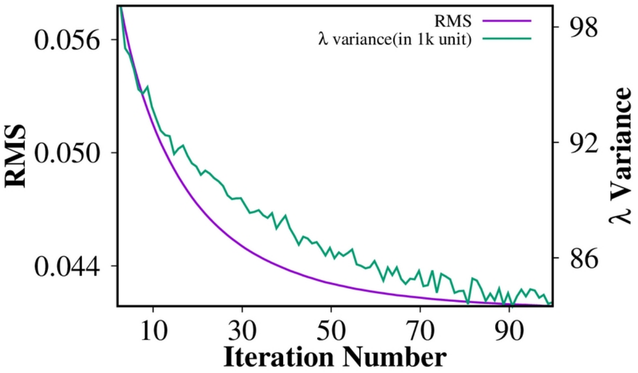

**Figure 2: Convergence of $\lambda$ values during MaxEnt inference for the nanog locus.** Root-mean-square (RMS; purple) and variance of $\lambda$ (green, in $k_BT$ units) are shown as a function of iteration number during the bias-relaxation cycles. Both metrics decrease and stabilize over time, indicating convergence of the inferred $\lambda$ parameters within $\sim$100 iterations.

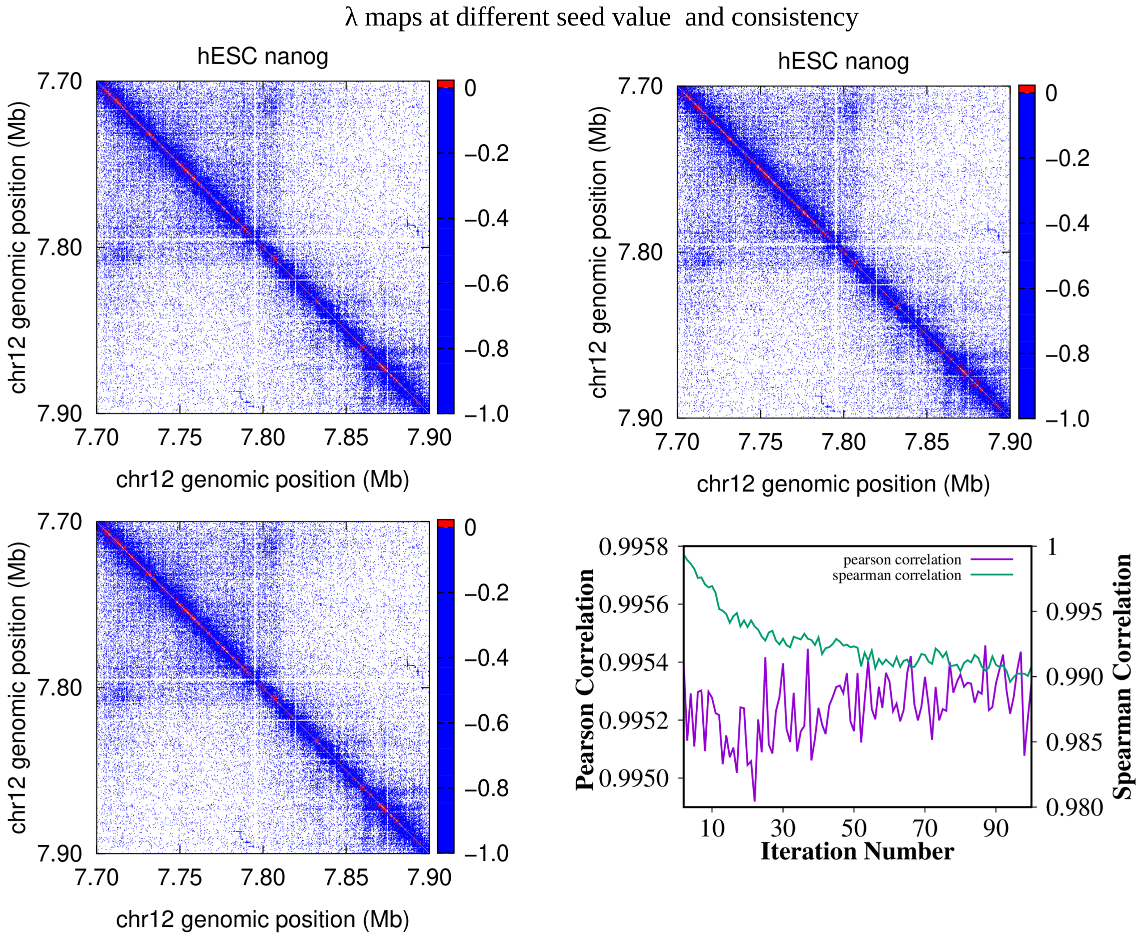

**Figure 3: Reproducibility of $\lambda$ maps across independent MaxEnt runs for the nanog locus in hESC.** The top and bottom-left panels show representative $\lambda$ maps obtained from different random seeds, demonstrating visually consistent spatial patterns. The bottom-right panel shows Pearson and Spearman correlations across iterations, indicating high reproducibility and stability of the inferred $\lambda$ parameters. For visualisation of $\lambda map$, $\lambda_{ij} valueswerenormalisedby \max(|\lambda|) withineachlocus$.

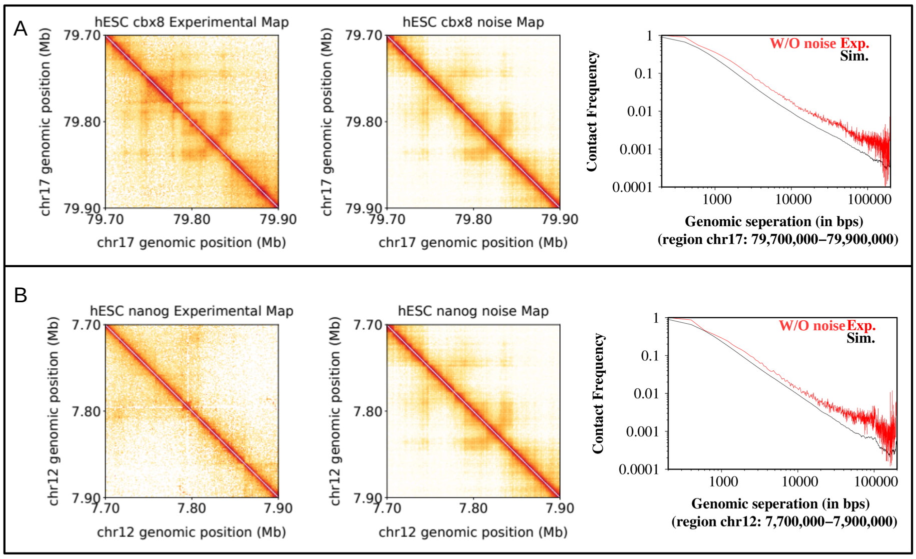

**Figure 4: Robustness of the MaxEnt inference to experimental noise in the input contact map. Top:** Original (left) and noise-perturbed (right) Micro-C contact maps for the hESC ppm1g and nanog loci ($\alpha = 0.5$). **Bottom:** MAXENT-simulated contact maps inferred from the noise-perturbed input compared with the original experimental map. Pearson and Spearman correlations between the noise-derived simulation and the original experimental map remain high (CBX8: $r = 0.924$, $\rho = 0.993$; nanog: $r = 0.881$, $\rho = 0.998$), confirming that the inference is not sensitive to moderate experimental perturbations.

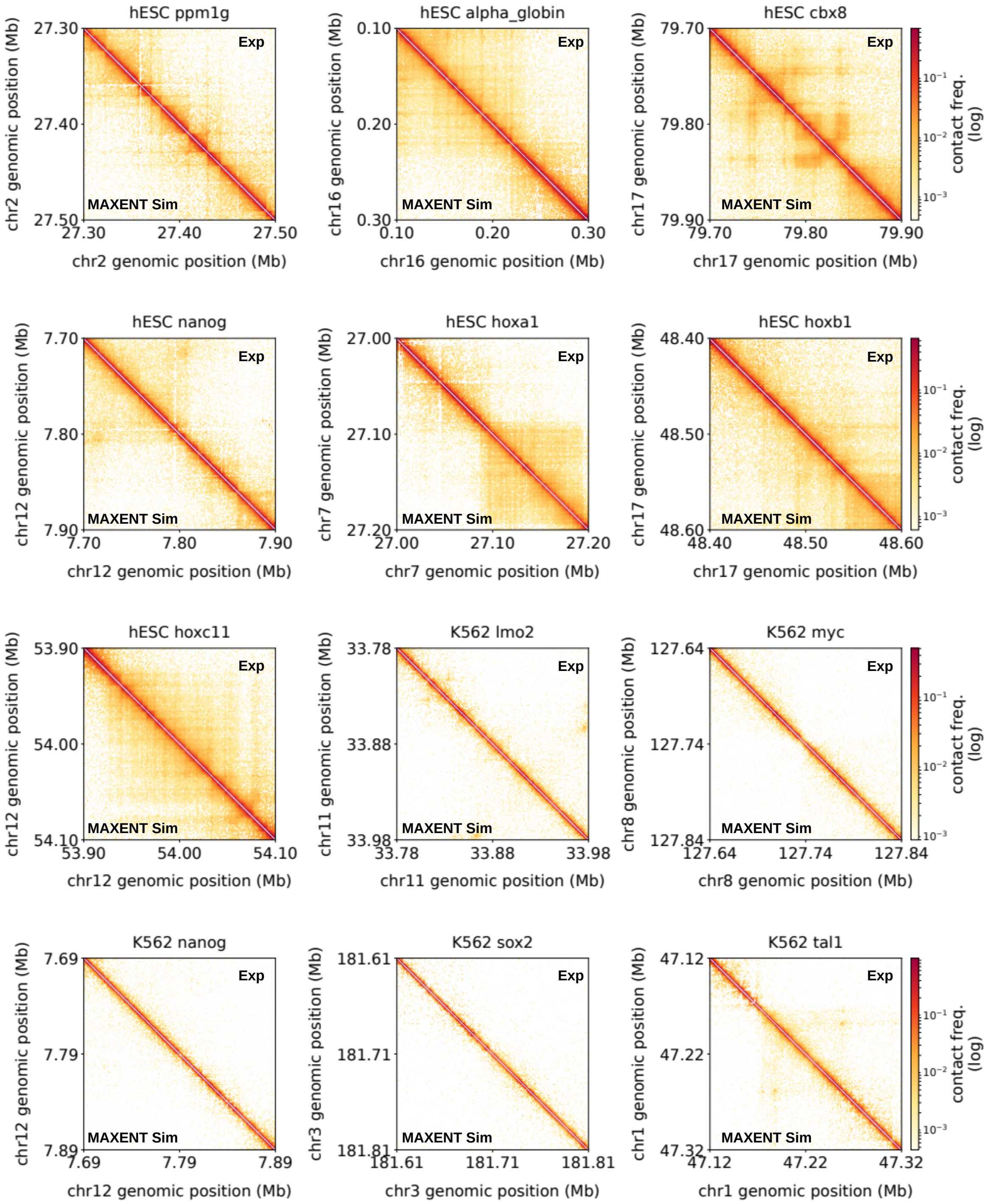

**Figure 5: Micro-C Contact map for loci in hESC and K562 cells** Experimental Micro-C contact maps showing chromatin interaction frequencies across 200 kb genomic regions in hESC and K562 cells.

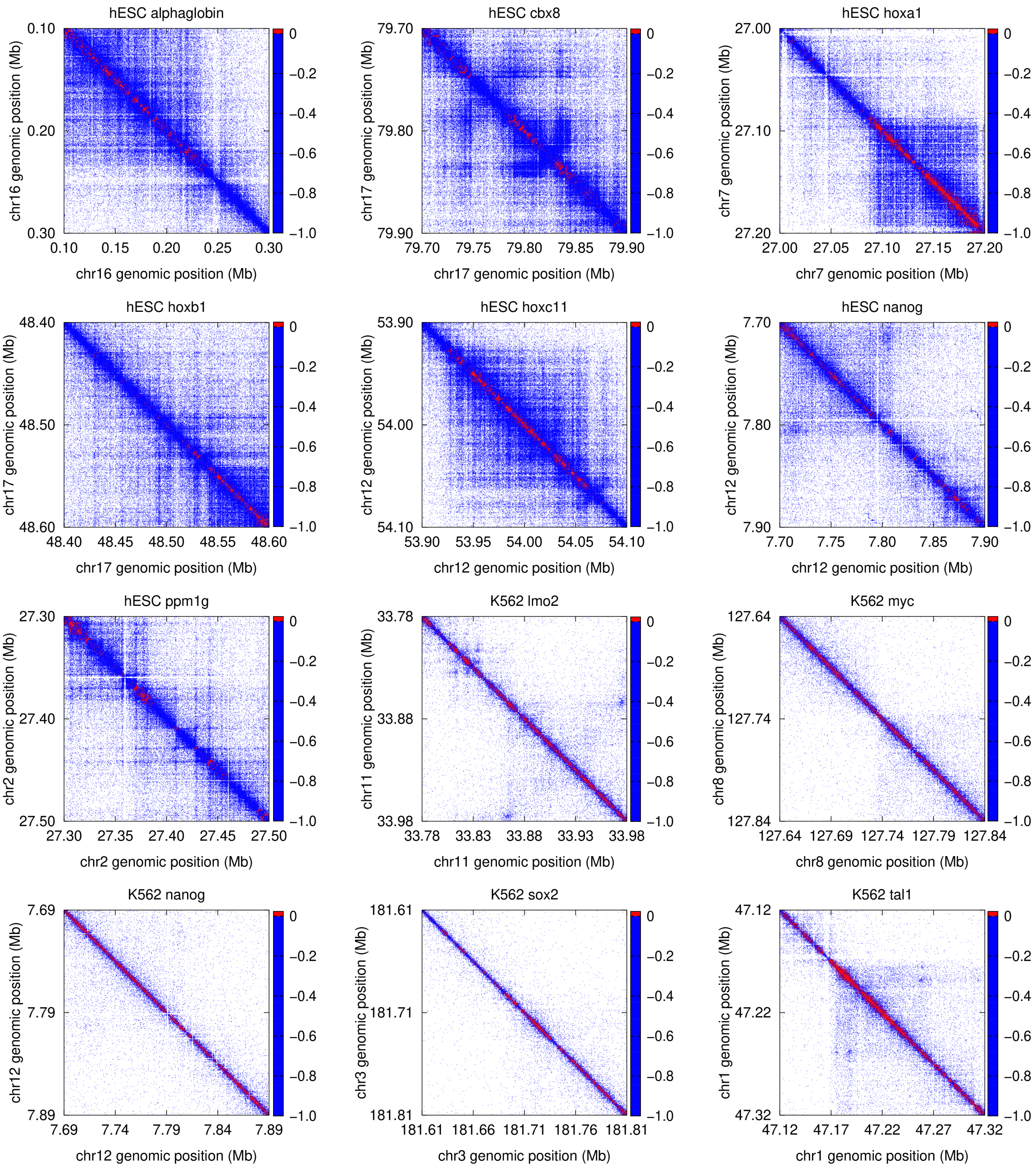

**Figure 6: Inferred $\lambda$ interaction landscapes across all 12 analysed loci.** For each locus (rows: hESC loci, top seven; K562 loci, bottom five), the experimental Micro-C contact map (left) and the corresponding inferred $\lambda$ map (right) are shown. The $\lambda$ maps are consistently structured across loci, exhibiting a sparse negative backbone concentrated near the diagonal and locus-specific long-range attractive couplings. The colour scale for $\lambda$ is the same in all panels. For visualisation of $\lambda map, \lambda_{ij} valueswerenormalisedby \max(|\lambda|) withineachlocus.$

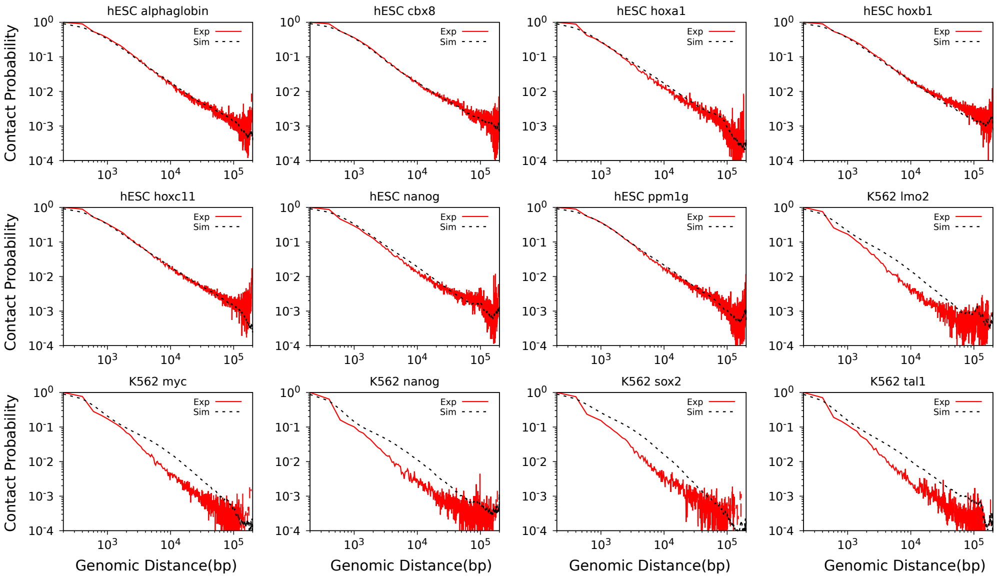

**Figure 7: Contact probability scaling $P(s)$ for MAXENT-simulated versus experimental contact maps across all analysed loci.** Contact probability $P(s)$ is plotted as a function of genomic separation $s$ (bp) for MAXENT-simulated (dashed lines) and experimental Micro-C (solid lines) maps over the range $10^2$-$10^5$ bp. Agreement extends over four decades of contact probability, confirming that the inferred $\lambda$ landscape captures both local and global structural scaling behaviour.

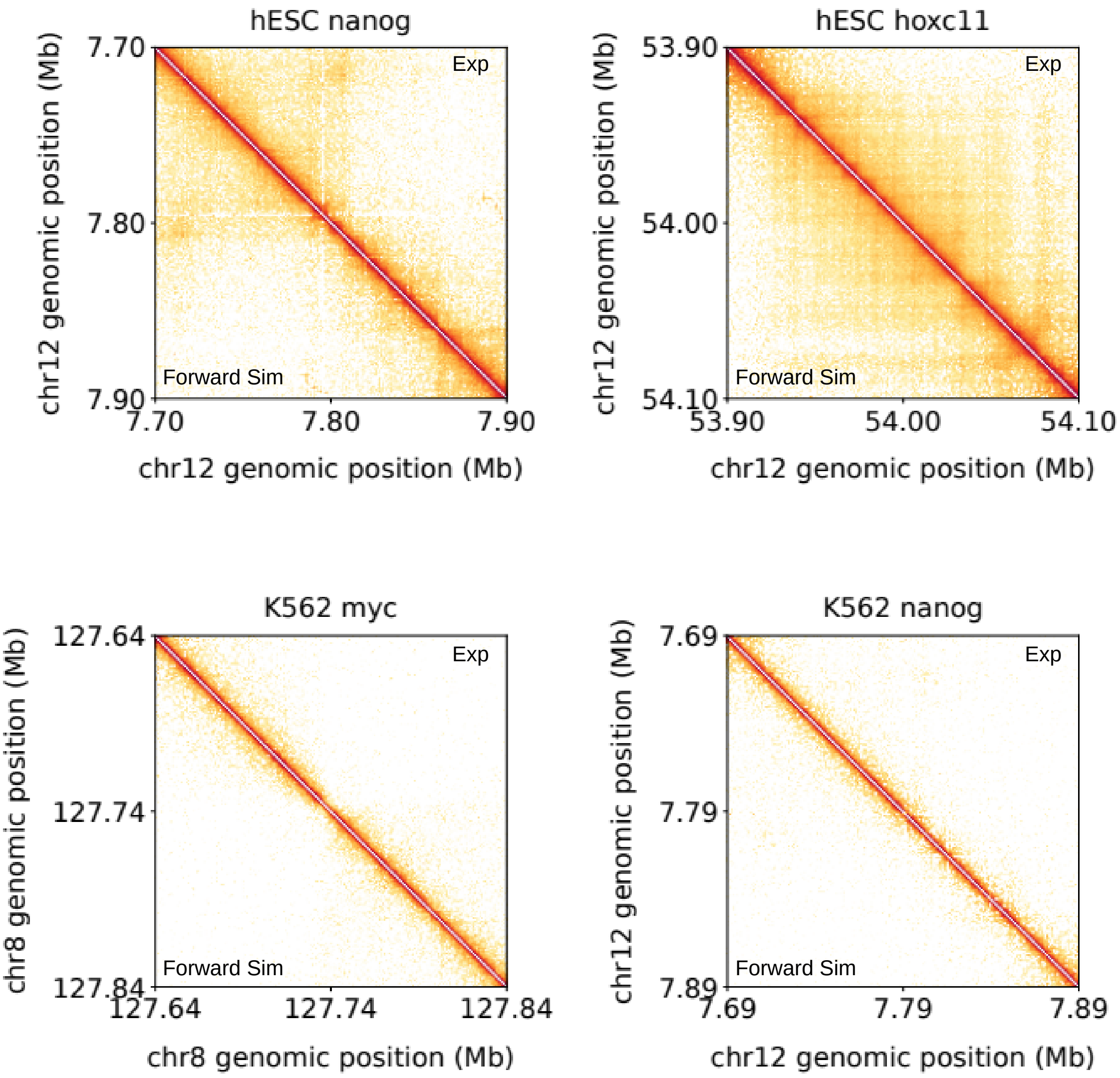

**Figure 8: Forward-simulation contact maps benchmarked against experimental Micro-C across 4 loci.** For each locus, the upper triangular portion shows the experimental Micro-C contact map and the lower triangular portion shows the corresponding forward-simulated map generated with the converged $\lambda$ landscape held fixed. Pearson and Spearman correlations are indicated for each locus. Key domain features (diagonal blocks, inter-domain boundaries) are visually reproduced in all cases.

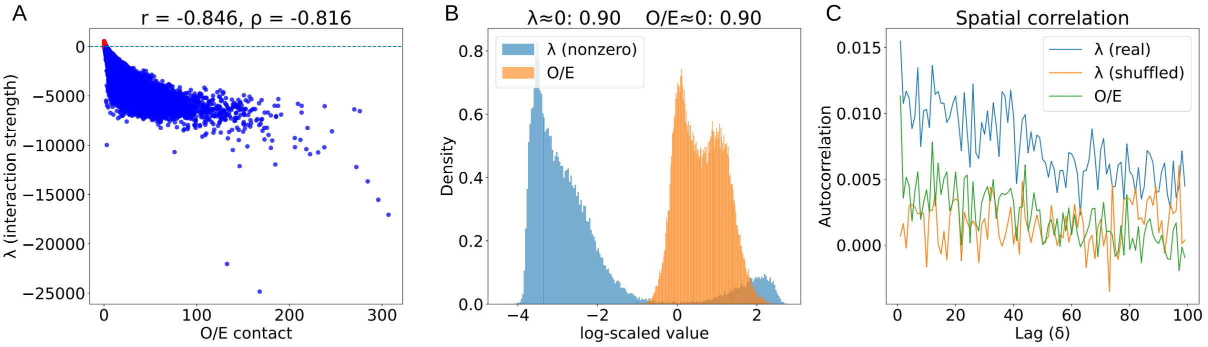

**Figure 9: Comparison of the $\lambda$ landscape with O/E-normalised contacts. Left:** Scatter plot of $|\lambda_{ij}|$ versus O/E-normalised contact enrichment $(C_{ij} - \langle C(s)\rangle)/\langle C(s)\rangle$, with Pearson ($r$) and Spearman ($\rho$) correlations indicated (hESC nanog locus). **Centre:** Distribution of non-zero $\lambda_{ij}$ entries (blue) compared with the non-zero distribution of the O/E map (orange), illustrating the greater sparsity of the $\lambda$ representation. **Right:** Spatial correlation of the real $\lambda$ map (blue) versus a distance-preserving shuffled control (grey) and the O/E map (orange). Coherent spatial correlations present in $\lambda$ are absent in the O/E map shuffled control, confirming that $\lambda$ encodes higher-order spatial organisation not accessible to O/E normalisation.

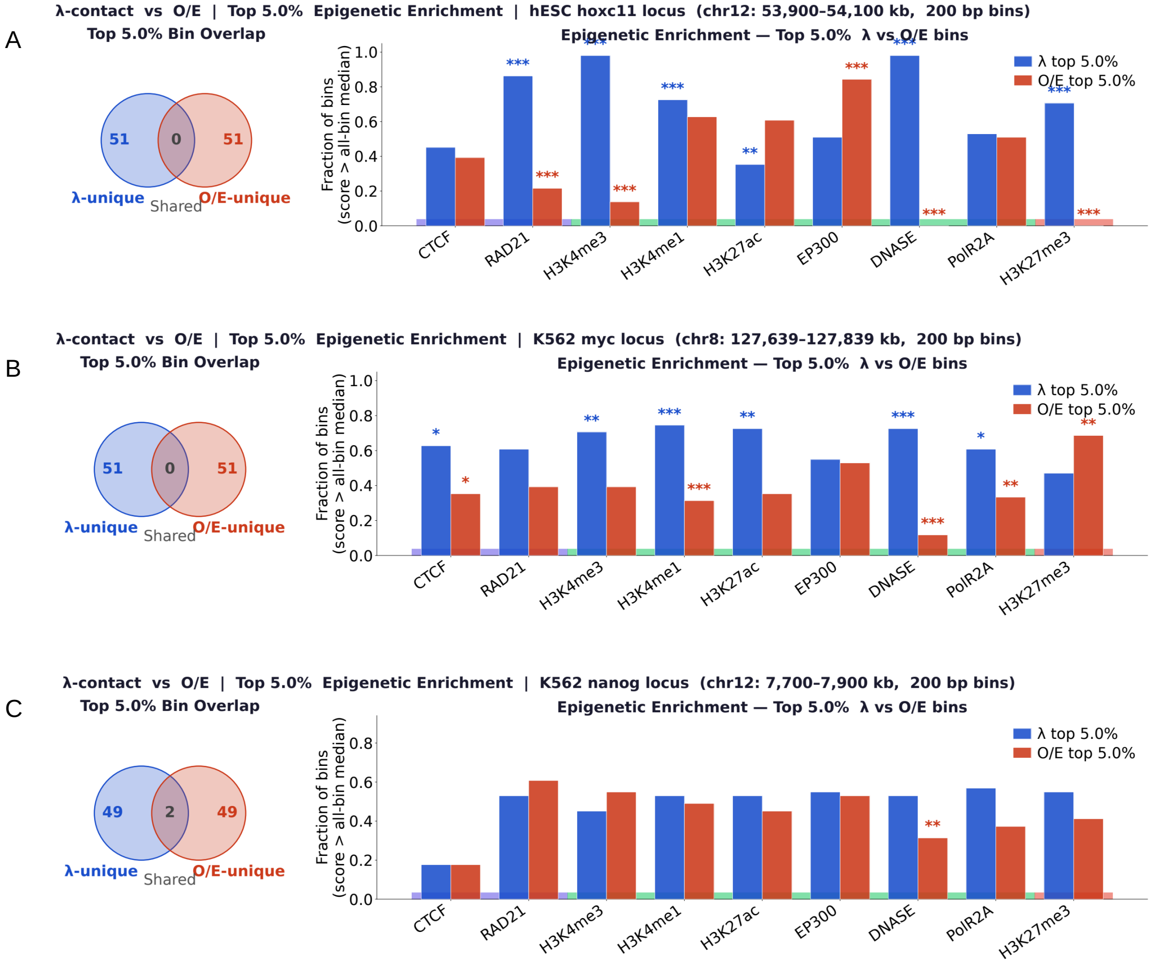

**Figure 10: The inferred landscape captures genomic features distinct from O/E-normalized contact frequency.** Left: Minimal overlap is observed between the top 5% of genomic bins ranked by $|\lambda|$ and O/E values, with only 1–2 shared bins among 50 bins in each set. Right: Epigenetic enrichment analysis reveals that $\lambda$-top bins are preferentially associated with architectural chromatin proteins and display enrichment levels comparable to or exceeding those of O/E-top bins across all tested chromatin marks. In contrast, O/E-top bins are primarily enriched for enhancer and promoter associated histone modifications. Statistical significance was assessed using the Mann–Whitney U-test (P $<$ 0.05–0.001).

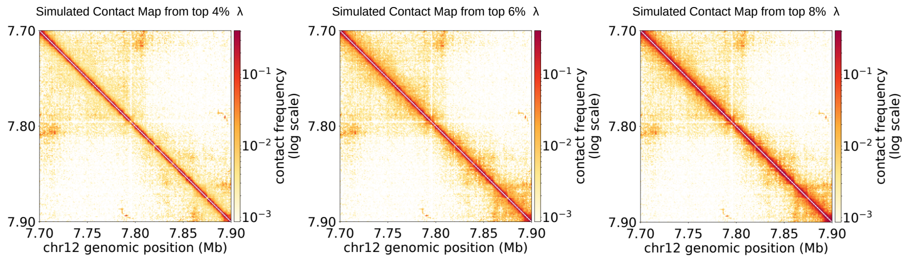

**Figure 11:** The top Forward-simulated contact maps reconstructed from progressively truncated $\lambda$ matrices containing only the top 4%, 6%, and 8% of interactions ranked by absolute magnitude. The resulting maps retain the principal domain organization and long-range contact patterns, indicating that a limited fraction of high-magnitude interactions is sufficient to recover the dominant features of chromatin architecture.

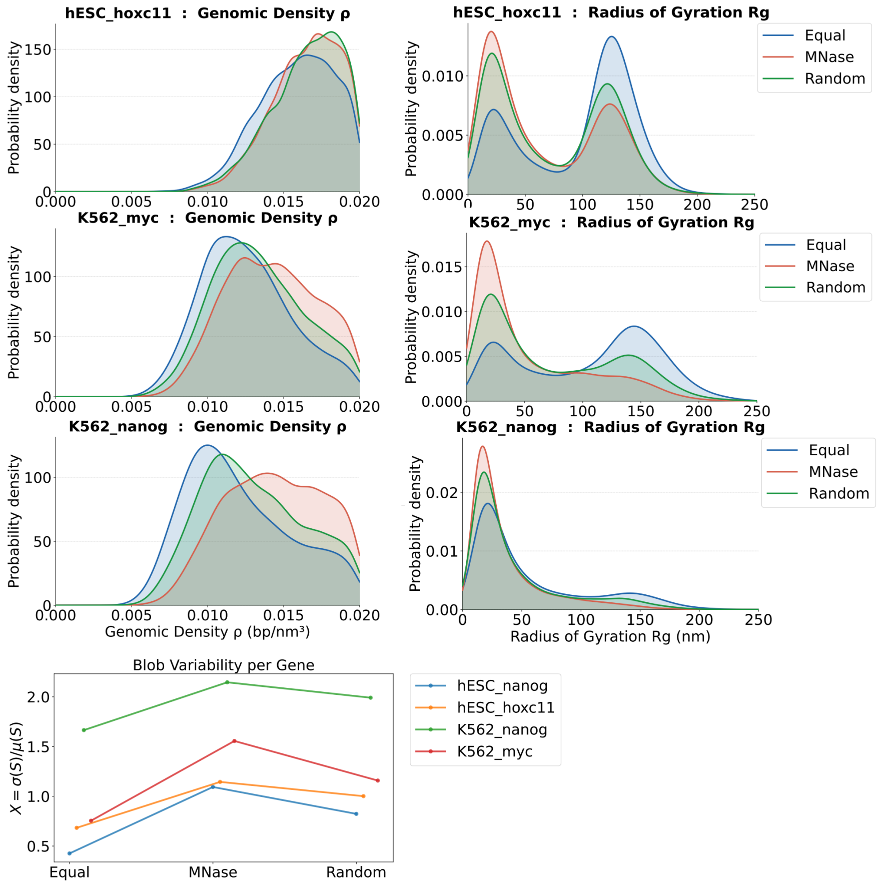

**Figure 12: Blob structural properties across all 12 analysed loci for the three nucleosome configurations.** Box plots of most probable blob size (bp), mean $R_g$ (nm), most probable $R_g$ (nm), and mean packing density (bp/nm$^3$) for MNase-seq (blue), randomised (orange), and uniformly spaced (green) nucleosome configurations, shown for each locus individually and as a cross-locus average. The MNase-seq configuration consistently yields the smallest, most compact, and densest blobs across all loci.

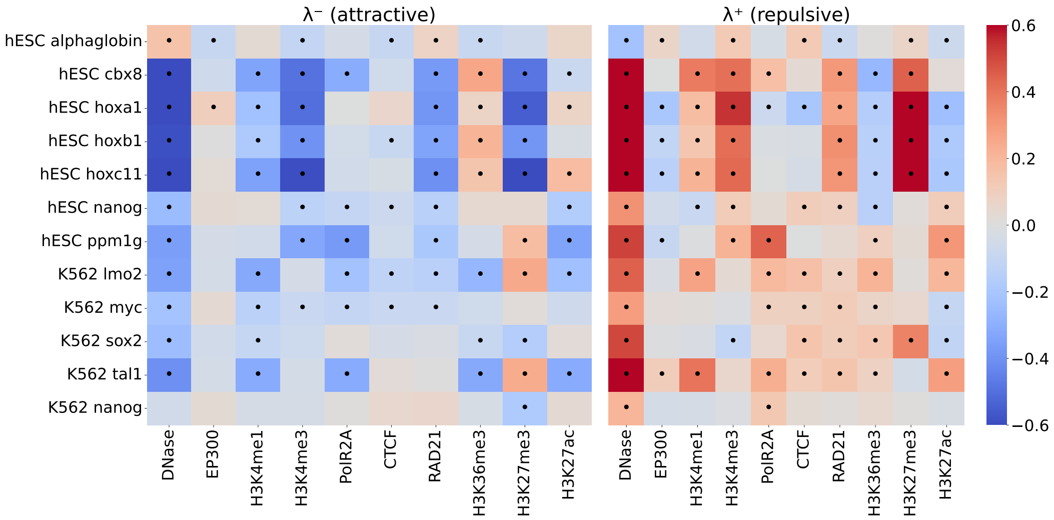

**Figure 13: Cross-locus epigenomic associations of $\lambda^-$ and $\lambda^+$ across all 12 analysed loci.** Spearman correlations between $\lambda^-$ (attractive, left) or $\lambda^+$ (repulsive, right) genomic profiles and ChIP-seq/ATAC-seq signal tracks. Marks are grouped as active (DNase, H3K27ac, H3K4me1, H3K4me3, EP300, POLR2A), architectural (CTCF, RAD21), and repressive (H3K36me3, H3K27me3). Dots indicate statistical significance after Benjamini–Hochberg correction ($q < 0.05$). The most reproducible cross-locus association is a positive correlation between $\lambda^+$ and DNase hypersensitivity, significant across the majority of loci. H3K27me3 association with $\lambda^+$ is restricted to Polycomb-regulated loci. $\lambda^-$ shows consistently negative correlation with DNase across loci.
